# Abnormal triaging of misfolded proteins by adult neuronal ceroid lipofuscinosis-associated CSPα mutants causes lipofuscin accumulation

**DOI:** 10.1101/2021.07.16.452648

**Authors:** Juhyung Lee, Yue Xu, Layla Saidi, Miao Xu, Konrad Zinsmaier, Yihong Ye

## Abstract

Mutations in *DNAJC5* (encoding the J domain-containing HSP70 co-chaperone CSPα) are associated with adult neuronal ceroid lipofuscinosis (ANCL), a dominant-inherited neurodegenerative disease featuring lysosome-derived <u>a</u>utofluorescent *<u>s</u>*torage <u>m</u>aterial (AFSM) termed lipofuscin. Functionally, CSPα has been implicated in chaperoning synaptic proteins and in misfolding-associated protein secretion (MAPS), but how CSPα dysfunction causes lipofuscinosis and neurodegeneration is unclear. Here we report two distinct protein quality control functions of CSPα at endolysosomes and perinuclear vesicles, respectively. We show that the endolysosome-associated CSPα promotes microautophagy of misfolded clients, but is dispensable for MAPS. By contrast, the perinuclear-localized CSPα, regulated by a previously unknown CSPα interactor named CD98hc, is critical for MAPS but unneeded for microautophagy. Importantly, these processes are coupled by CSPα in a J-domain regulated manner. Uncoupling these two processes, as seen in cells lacking CD98hc or expressing ANCL-associated CSPα mutants, generates CSPα-containing AFSMs resembling NCL patient-derived lipofuscin, and also induces neurodegeneration in a *Drosophila* ANCL model. These findings suggest that blocking MAPS while allowing CSPα-mediated microautophagy disrupts lysosome homeostasis, causing CSPα-associated lipofuscinosis and neurodegeneration.

---

Neuronal Ceroid Lipofuscinosis (NCL) refers to a family of genetically inherited neurodegenerative lysosomal storage diseases that are associated with an excessive accumulation of lipopigments (lipofuscin), which occurs in both neurons and non-neuronal tissues ^1, 2^. A key feature of lipofuscin is its autofluorescence, which allows detection by fluorescence microscopy ^3^. These diseases can occur in either infants, juveniles, or adults. The infantile and juvenile variants of NCL (INCL and JNCL) are often more severe, associated with progressive vision loss, seizure, and brain death. The adult variant (ANCL also named CLN4) on the other hand has milder symptoms. Nevertheless, after diagnosis, ANCL patients usually die after 10 years ^4^.

Mutations in a collection of CLN genes (Ceroid Lipofuscinosis Neuronal) have been linked to various NCL variants ^4–6^. Some of these genes encode proteins essential for lysosomal functions such as enzymes mediating lysosomal degradation (e.g. PPT1 and CTSD) ^4^ or regulators governing the trafficking of lysosome resident proteins (e.g. CLN6 and CLN8) ^7, 8^. These observations prompted the idea that lipofuscin accumulation and neurodegeneration in NCL might be a result of disrupted lysosome homeostasis. However, how CLN mutations cause lipofuscin build-up and neurodegeneration has remained largely elusive.

The ANCL is caused by dominant mutations in the gene encoding cysteine string protein-α (CSPα, also named DNAJC5), which is a vesicle-associated protein. CSPα features three conserved domains: an HSC70-binding J-domain near the N-terminus, a central cysteine string (CS) domain, and a linker (LN) domain sandwiched between the J domain and the CS domain ^9^. The cysteine residues in the CS domain are mostly palmitoylated, which regulates CSPα membrane association and trafficking ^10, 11^. In non-neuronal cells, CSPα is found on the plasma membrane and in association with endolysosomes ^12, 13^. However, in neurons, it is mostly associated with synaptic vesicles ^14–16^. As an Hsc70 co-chaperone, CSPα can stimulate the ATPase activity of Hsc70 and Hsp70 in vitro ^17, 18^. In neurons, it can function in conjunction with both chaperones to control the folding and trafficking of synaptic proteins such as SNAP25 and Dynamin ^19, 20^, which in term regulate a variety of cellular processes including calcium homeostasis ^21–24^, membrane fusion ^25, 26^, neurotransmitter release ^26–28^ and synapse stability ^29–31^. Moreover, recent studies have also implicated CSPα in the unconventional secretion of misfolded cytosolic proteins via a process named misfolding-associated protein secretion (MAPS) ^12, 32, 33^. Since the MAPS activity of CSPα was tightly linked to its endolysosome association, it was proposed that CSPα might chaperone misfolded proteins to endolysosomes for their secretion to the cell exterior^32^.

Two ANCL-associated mutations have been identified ^34–36^, which are located next to each other in the CS domain, close to the interface of the upstream LN domain whose function is unclear. Recent studies suggest that these mutations reduce CSPα palmitoylation while increasing its aggregation propensity ^13, 37–39^. These mutations also cause mislocalization of the protein in cells ^38^. Accordingly, ANCL-causing *CSPα* mutations are thought to reduce CSPα chaperoning function ^39^. However, this notion does not explain why cells carrying *CSPα* mutations accumulate lysosome-derived lipopigments.

In this study, we found that CSPα used a J-domain independent activity to couple two protein quality control (PQC) processes: ESCRT-dependent microautophagy and misfolding-associated protein secretion (MAPS). We identified CD98hc as a CSPα interactor critical for MAPS, but dispensable for microautophagy. Importantly, CD98hc depletion or expression of ACNL-associated CSPα mutants uncouple the two PQC processes, resulting in lipofuscinosis and neurodegeneration.

## Results

### J-domain independent translocation of CSPα into the lumen of endolysosomes

Since CSPα was detected on lysosomes and in conditioned medium ^12^, we asked whether it could enter the lumen of endolysosomes prior to its release into the cell exterior. To this end, we established a Keima-based fluorescence assay. Keima is a monomeric dual-excitable fluorescence protein (λ^em^max ∼620 nm) with maximum excitation at 440 nm under neutral pH (6.0-8.0) or at 550 nm in an acidic environment (pH<6.0) (Fig. 1a) ^40^. When fused to a cargo, Keima allows quantitative measurement of cargo trafficking to the acidic endolysosomes by fluorescence microscopy or flow cytometry ^41^.

**Figure 1.**
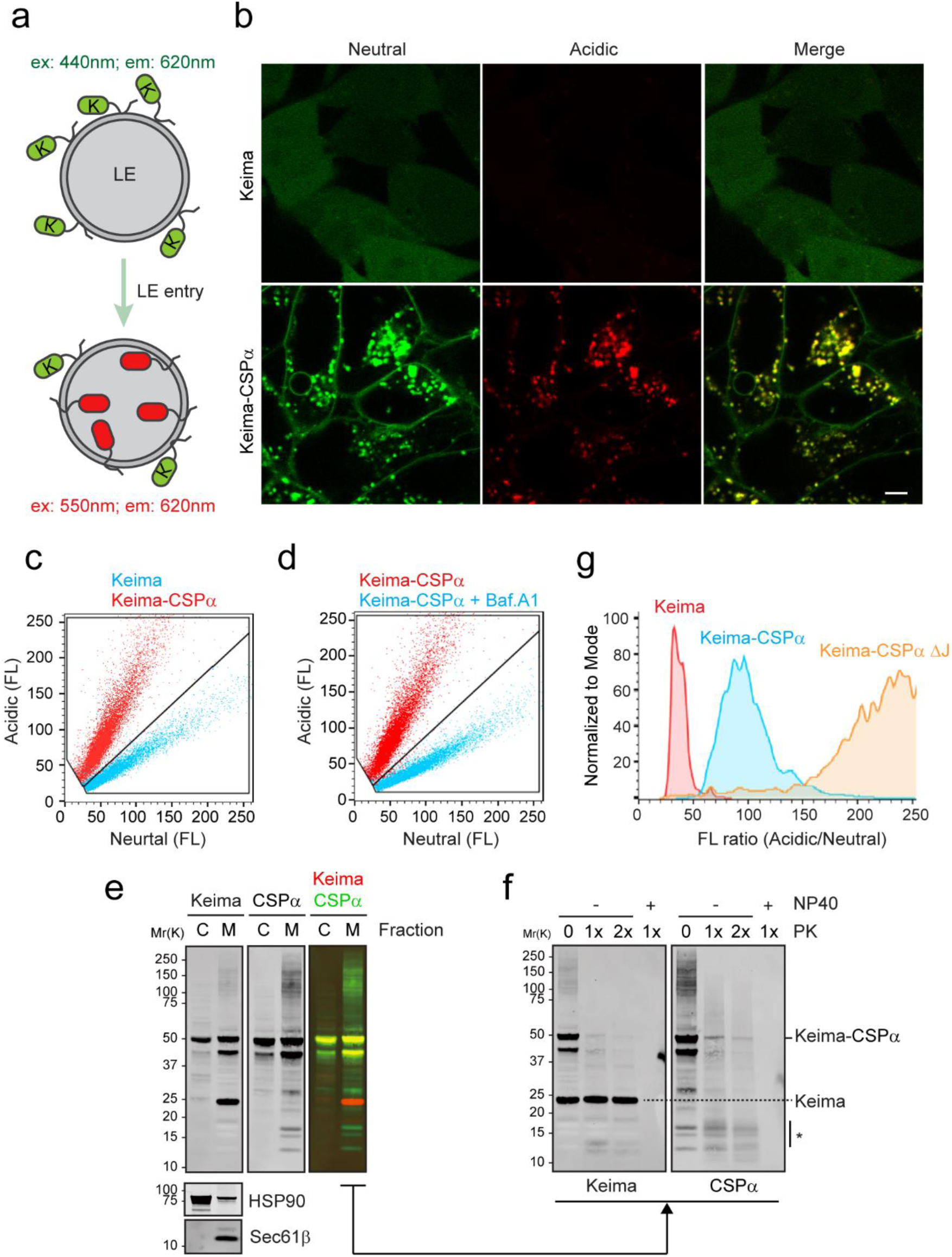
Translocation of CSPα into the lumen of endolysosomes. **a,** The Keima-based endolysosome translocation assay. **b, c,** Keima-CSPα but not Keima is efficiently translocated into endolysosomes. HEK293T cells stably expressing Keima or Keima-CSPα were analyzed by confocal microscopy (b) or flow cytometry (c). Scale bar, 5 μm. **d,** Flow cytometry analysis of Keima-CSPα cells before and after bafilomycin A1 (200 nM, 2 h) treatment. **e, f,** Membrane fractionation and protease protection demonstrate the translocation of CSPα into endolysosomes. e, The membrane (M) and cytosol (C) fractions from Keima-CSPα cells were analyzed by immunoblotting with the indicated antibodies. f, Immunoblotting analysis of the membrane fraction from e after treatment with increased concentrations of proteinase K in the absence (-) or presence (+) of 0.1% NP40. Asterisk indicates protease-resistant CSPα fragments. **g,** Deletion of the J-domain stimulates CSPα endolysosome translocation. The ratio of acidic vs. neutral fluorescence signals in cells stably expressing the indicated Keima proteins was analyzed by FACS.

Confocal microscopy showed that HEK293T cells stably expressing Keima displayed only neutral cytoplasmic fluorescence. By contrast, in Keima-CSPα expressing cells, we not only detected a neutral Keima-CSPα signal on the plasma membrane but also observed Keima-CSPα on intracellular vesicles by excitation at both 440 nm and 550 nm (Fig. 1b). Thus, these vesicles are endolysosomes containing a fraction of CSPα on the surface and in the lumen. Consistently, flow cytometry revealed a much higher acidic fluorescence signal in Keima-CSPα cells than in Keima cells (Fig. 1c). As anticipated, treatment of bafilomycin A1 (Baf. A1), a lysosome acidification blocker, dramatically reduced the acidic Keima-CSPα signal but increased the neutral Keima-CSPα signal (Fig. 1d). These findings demonstrate that CSPα can enter the lumen of endolysosomes.

We used membrane fractionation and a protease protection assay to further monitor the endolysosome translocation of Keima-CSPα. We detected full-length Keima-CSPα in both the cytosol and membrane fractions, and several proteolyzed species in the membrane fraction only (Fig. 1e). A 25kD cleavage product was detected by Keima antibodies, whereas several lower molecular weight species were detected only by CSPα antibodies. Thus, Keima-CSPα is cleaved in endolysosomes by proteases that spared Keima, consistent with Keima being a hydrolase-resistant protein ^41^. Treating membranes with proteinase K caused rapid degradation of most full-length Keima-CSPα as well as a 42kD species. By contrast, the 25kD Keima protein, a small fraction of full-length CSPα, and the low molecular weight CSPα cleavage products were protected unless the detergent NP40 was added (Fig. 1f). These results further confirm the endolysosomal translocation of Keima-CSPα.

To test if the endolysosome translocation of CSPα requires the J-domain, we measured the ratio of the acidic and neutral fluorescence signal in cells stably expressing either WT Keima-CSPα or a Keima-CSPα mutant lacking the J-domain (ΔJ). Intriguingly, deleting the J-domain further enhanced the endolysosome translocation of CSPα (Fig. 1g). The effect of J-domain deletion on CSPα endolysosome translocation was likely achieved via a gain-of-function activity because this mutant was also more active in promoting MAPS (see below).

### CSPα promotes endolysosomal translocation of misfolded proteins via microautophagy

To test whether CSPα may chaperone misfolded proteins to the lumen of endolysosomes, we used a truncated GFP protein (GFP1-10) as a misfolded model substrate because we previously detected the association of a small fraction of mCherry-tagged GFP1-10 with endolysosomes using a photobleaching-based live-cell imaging assay ^32^. In the control cells expressing only mCh-GFP1-10, a few mCherry-positive vesicles were detected after photobleaching. Co-expression of CSPα increased the endolysosome association of mCh-GFP1-10, which was further enhanced in CSPα ΔJ-expressing cells (Fig. 2a panels 3, 2 vs. 1, Fig. 2b). As in CSPα WT cells, the mCh-GFP1-10-containing vesicles induced by CSPα ΔJ also contain CSPα ΔJ and the late endosome protein Rab9 (Fig. 2a, panels 4-9). Taking together, this suggests that CSPα recruits mCh-GFP1-10 to endolysosomes independent of its J-domain.

**Figure 2.**
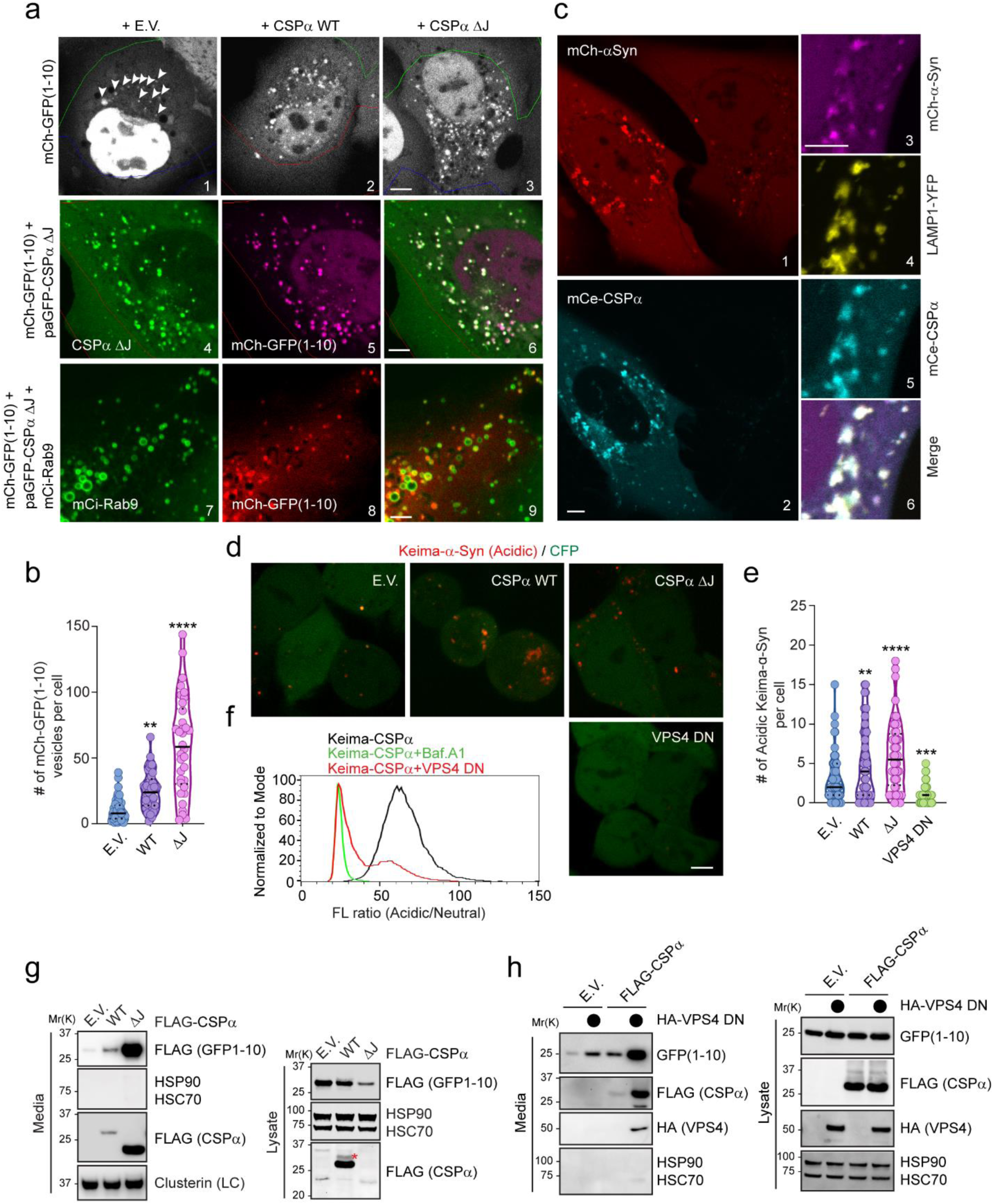
CSPα-mediated microautophagy is dispensable for MAPS. **a-c,** CSPα promotes the association of misfolded proteins with endolysosomes. a, Panels 1-3, COS-7 cells transfected with mCh-GFP1-10 together with either empty vector (EV) or photoactivatable GFP-tagged CSPα WT or CSPα ΔJ were subject to repeated photobleaching in the indicated area, which reveals vesicle-associated mCh-GFP1-10 (arrowheads). The number of mCh-GFP1-10-containing vesicles in individual cells are shown in b. **, *p* < 0.005 (*p* = 0.0023); ****, *p* < 0.0001 by one-way ANOVA plus Dunnett’s test; *n* = 50, 43, and 34 cells respectively. Panels 4-6 show the co-localization of mCh-GFP1-10 with paGFP-CSPα ΔJ. Panels 7-9, cells transfected with mCh-GFP1-10, CSPα ΔJ, and mCi-Rab9 show the co-localization of mCh-GFP1-10 with Rab9. c, U2OS cells stably expressing mCh-α-Syn were transfected with mCerulean (mCe)-CSPα and LAMP1-YFP and imaged. The right panels show enlarged views from another cell. Scale bars, 5 μm. **d, e,** CSPα promotes the endolysosome translocation of Keima-α-Syn. Keima-α-Syn stable HEK293T cells transfected with the indicated plasmids together with a CFP-expressing plasmid were imaged. CFP serves as a control for transfected cells. Scale bar, 5 μm. e shows the number of acidic Keima-α-Syn-containing vesicles in individual cells in d. **, *p* < 0.005 (*p* = 0.0048); ***, *p* < 0.0005 (*p* = 0.0003); ****, *p* < 0.0001 by one-way ANOVA plus Dunnett’s test. *N* = 63, 46, 48, and 77 cells, respectively. **f,** VPS4 DN inhibits the endolysosomal translocation of Keima-CSPα. Keima-CSPα stable cells were treated with bafilomycin A1 or transfected with either an empty vector or VPS4 DN (E228Q)-expressing plasmid, and then analyzed by FACS. **g, h,** CSPα-mediated secretion of misfolded proteins is independent of endolysosomal translocation. Conditioned medium and cell lysates from HEK293T cells transfected with FLAG-GFP1-10 together with the indicated plasmids were analyzed by immunoblotting. LC, loading control.

We performed a similar experiment using α-Synuclein (α-Syn), a misfolding-prone protein associated with Parkinson’s disease ^42^. U2OS cells stably expressing mCherry-tagged human α-Syn showed mainly cytoplasmic mCh-α-Syn signal, but a few α-Syn-containing vesicles could be detected even without photobleaching. CSPα expression significantly stimulated α-Syn membrane association, resulting in many bright mCh-α-Syn positive punctae (Fig. 2c, panels 1, 2). As expected, vesicle-associated α-Syn was co-localized with CSPα and the lysosomal marker LAMP1 (Fig. 2c, panels 3-6). Thus, CSPα promotes the association of misfolded proteins with endolysosomes.

We next tested whether CSPα could promote the translocation of misfolded proteins into endolysosomes using cells stably expressing Keima-α-Syn. Confocal microscopy detected Keima-α-Syn mostly as cytoplasmic neutral fluorescence with a few acidic punctae, indicating that Keima-α-Syn is mostly localized in the cytoplasm. When CSPα WT or ΔJ was co-expressed, however, the number of acidic Keima-α-Syn punctae was significantly increased (Fig. 2d, e).

Microautophagy is a major known mechanism that translocates endolysosome-associated proteins to the lumen using the ESCRT complexes via multivesicular body formation ^43–46^. To test whether microautophagy is involved in CSPα-induced endolysosomal translocation of misfolded proteins, we ectopically expressed a dominant-negative (DN) VPS4 mutant (E228Q), which blocked ESCRT-mediated microautophagy ^47^. As expected, VPS4 DN expression abolished the acidic Keima-α-Syn punctae (Fig. 2d, e) and reduced the acidic Keima-CSPα fluorescence as shown by flow cytometry (Fig. 2f). Lysotracker staining showed that VPS4 DN did not affect the lysosomal pH (Supplementary Fig. 1a). Collectively, these results suggest that the endolysosomal translocation of CSPα and its clients is mediated by ESCRT-dependent microautophagy.

### Endolysosomal translocation of misfolded proteins is dispensable for MAPS

We next asked whether CSPα-mediated microautophagy is involved in MAPS. To this end, conditioned medium and cell lysate from HEH293T cells transfected with the MAPS substrate GFP1-10 ^32^ were analyzed by immunoblotting, which detected a fraction of GFP1-10 but not abundant cytosolic proteins such as Hsc70 and Hsp90 in the medium (Fig. 2g; Supplementary Fig. 1b). CSPα overexpression enhanced GFP1-10 secretion, consistent with previous reports ^12, 33^. Strikingly, the MAPS-stimulating activity of CSPα ΔJ was almost 10-fold higher than that of WT CSPα (Fig. 2g, Supplementary Fig. 1b). A similar observation was made for α-Syn (Supplementary Fig. 1c). Notably, a fraction of WT CSPα was also secreted, and the secretion of CSPα ΔJ was much higher than that of WT CSPα (Fig. 2g). The correlation between the endolysosomal translocation of CSPα and its MAPS-stimulating activity appears to suggest that endolysosomes may be a secretory compartment for MAPS. However, when we examined the secretion of GFP1-10 in the presence of VPS4 DN, surprisingly, VPS4 DN dramatically increased GFP1-10 secretion under both basal and CSPα-overexpressing conditions (Fig. 2h). These results suggest that CSPα-mediated microautophagy and MAPS are two parallel but functionally coupled processes, as both are regulated by the CSPα J-domain.

### ANCL mutations inhibit MAPS without affecting CSPα endolysosomal translocation

To test whether the above-mentioned CSPα functions were affected by ANCL-associated mutations, we first tested whether CSPα L115R and L116Δ mutants still promote endolysosome association of misfolded α-Syn using mCh-α-Syn-expressing cells. Like WT CSPα, both mCitrine (mCi)-tagged CSPα L115R and CSPα L116Δ stimulated endolysosome association of mCh-α-Syn (Fig. 3a, b). When the endolysosome translocation of Keima-α-Syn was analyzed, the disease-associated mutants were as active as WT CSPα in stimulating acidic Keima-α-Syn positive punctae (Fig. 3c, d). Importantly, when cells were transiently transfected with Keima-tagged WT CSPα or the ANCL-associated mutants, both Keima-CSPα L115R and L116Δ were translocated into endolysosomes more efficiently than WT CSPα (Fig. 3e, f). These results suggest that the ANCL mutations do not affect the microautophagy-promoting activity of CSPα.

**Figure 3.**
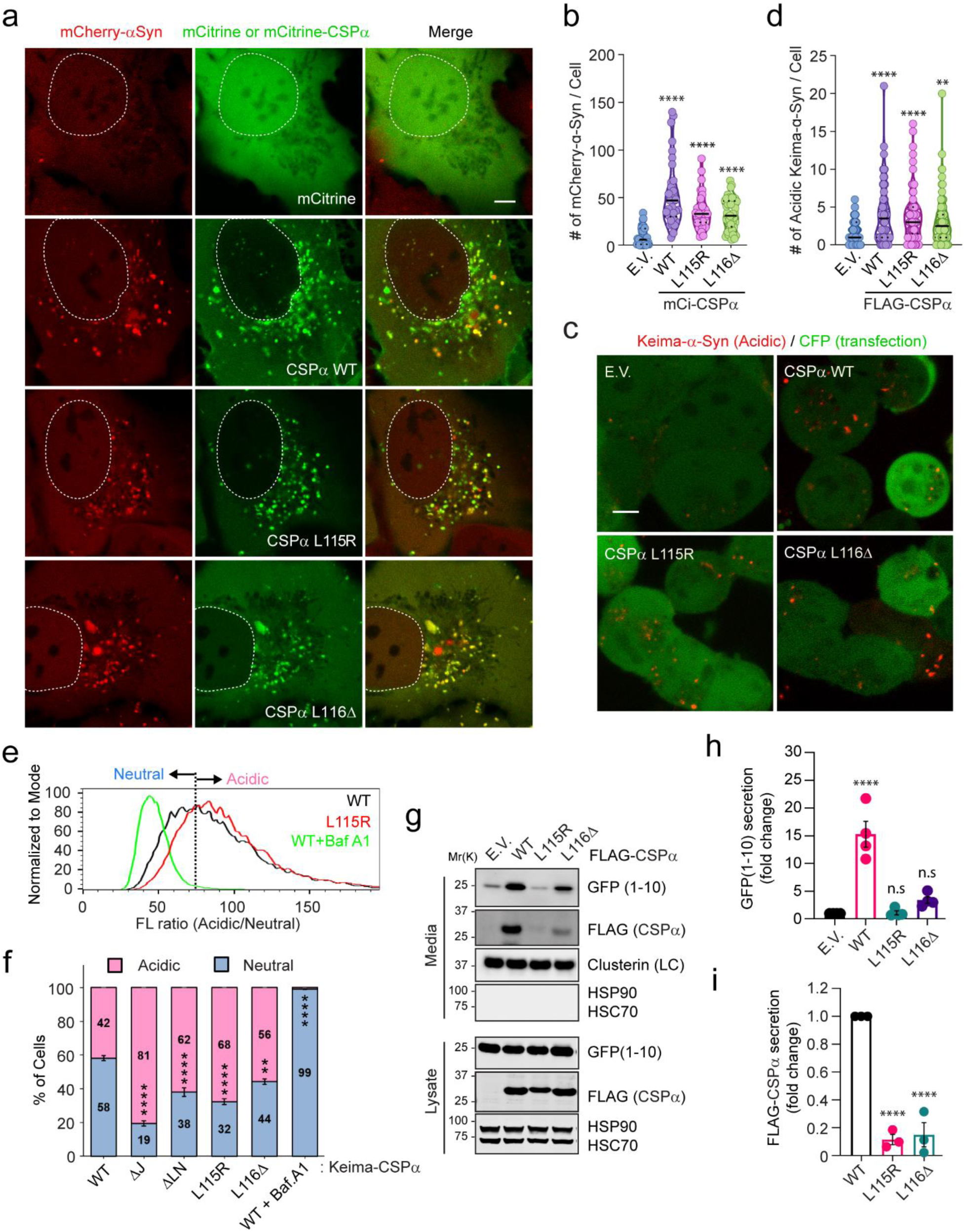
ANCL-associated CSPα mutants are defective in MAPS but capable of translocating substrates into endolysosomes. **a, b,** The CSPα L115R and L116Δ mutants promote the association of mCh-α-Syn with endolysosomes similarly as WT CSPα. Representative images of mCh-α-Syn-expressing U2OS cells transfected with the indicated plasmids. Scale bar, 5 μm. b, Quantification of mCh-α-Syn-containing vesicles in individual cells in a. ****, *p* < 0.0001 by one-way ANOVA plus Dunnett’s test; *n* = 37, 33, 31, and 45 cells, respectively. **c, d,** The CSPα L115R and L116Δ mutants can still promote the translocation of Keima-α-Syn into endolysosomes. c, Representative images of Keima-α-Syn cells transfected with the indicated plasmids together with CFP. Scale bar, 5 μm. d, Quantification of mCh-α-Syn-containing vesicles in individual cells in c. **, *p* < 0.005 (*p* = 0.0026); ****, *p* < 0.0001 by one-way ANOVA plus Dunnett’s test; *n* = 81, 60, 69 and 88 cells, respectively. **e, f,** The ANCL-associated CSPα mutants are translocated into endolysosomal lumen more efficiently than WT CSPα. e, Representative histograms show the ratio of acidic vs. neutral fluorescence intensity in cells transfected with the indicated plasmids. As a control, cells expressing WT CSPα were treated with Bafilomycin A1 (200 nM) for 2 h. The line indicates the threshold by which cell populations were assigned and analyzed in f. f, Quantification of cell populations as indicated in e in cells transfected with the indicated plasmids. error bars indicate mean ± s.e.m.; **, *p* < 0.005 (*p* = 0.0016); ****, *p* < 0.0001 by one-way ANOVA plus Dunnett’s test; *n*=3 experiments. **g-i,** The ANCL-associated CSPα mutants are defective in MAPS. Conditioned medium and cell lysates from cells transfected with GFP1-10 together with the indicated plasmids were analyzed by immunoblotting. LC, loading control. The graph in h shows the quantification of the secreted GFP1-10 normalized by GFP1-10 in cell lysates. Error bars indicate mean ± s.e.m. from *n*=4 independent experiments. The graph in i shows the secretion of CSPα from *n*=3 independent experiments; ****, *p* < 0.0001 by one-way ANOVA plus Dunnett’s test; n.s. not significant.

We next examined whether the ANCL mutations alter the CSPα function in MAPS. Interestingly, unlike WT CSPα, both the CSPα L115R and CSPα L116Δ mutants failed to promote GFP1-10 secretion (Fig. 3g, h). Likewise, the ANCL-associated CSPα mutants also failed to induce the secretion of α-Syn (Supplementary Fig.1d). Importantly, compared to WT CSPα, the secretion of the CSPα L115R and L116Δ mutants was also significantly reduced (Fig. 3g, i). These findings suggest that the CS domain-dependent palmitoylation plays an important role in MAPS, although it is non-essential for microautophagy.

### A linker domain targets CSPα to LAMP1-negative perinuclear vesicles

We next wished to identify the secretory compartment through which CSPα promotes MAPS. To this end, we re-characterized the subcellular localization of CSPα using HEK293T cells bearing a GFP at the carboxyl terminus of endogenous CSPα by 3D confocal microscopy, which revealed the presence of a fraction of CSPα in perinuclear vesicles that were largely negative for LAMP1 or the lysosome-specific dye Lysotracker (Supplementary Fig. 2a, b). A fraction of mCi-CSPα expressed in U2OS cells was also present in a LAMP1-negative perinuclear compartment (Fig. 4a, panels 1-6). By contrast, the ANCL-associated mutants were largely absent from this compartment. As a result, both CSPα L115R and L116Δ showed increased co-localization with LAMP1 (Fig. 4a panels 7-12, and Fig. 4b).

**Figure 4.**
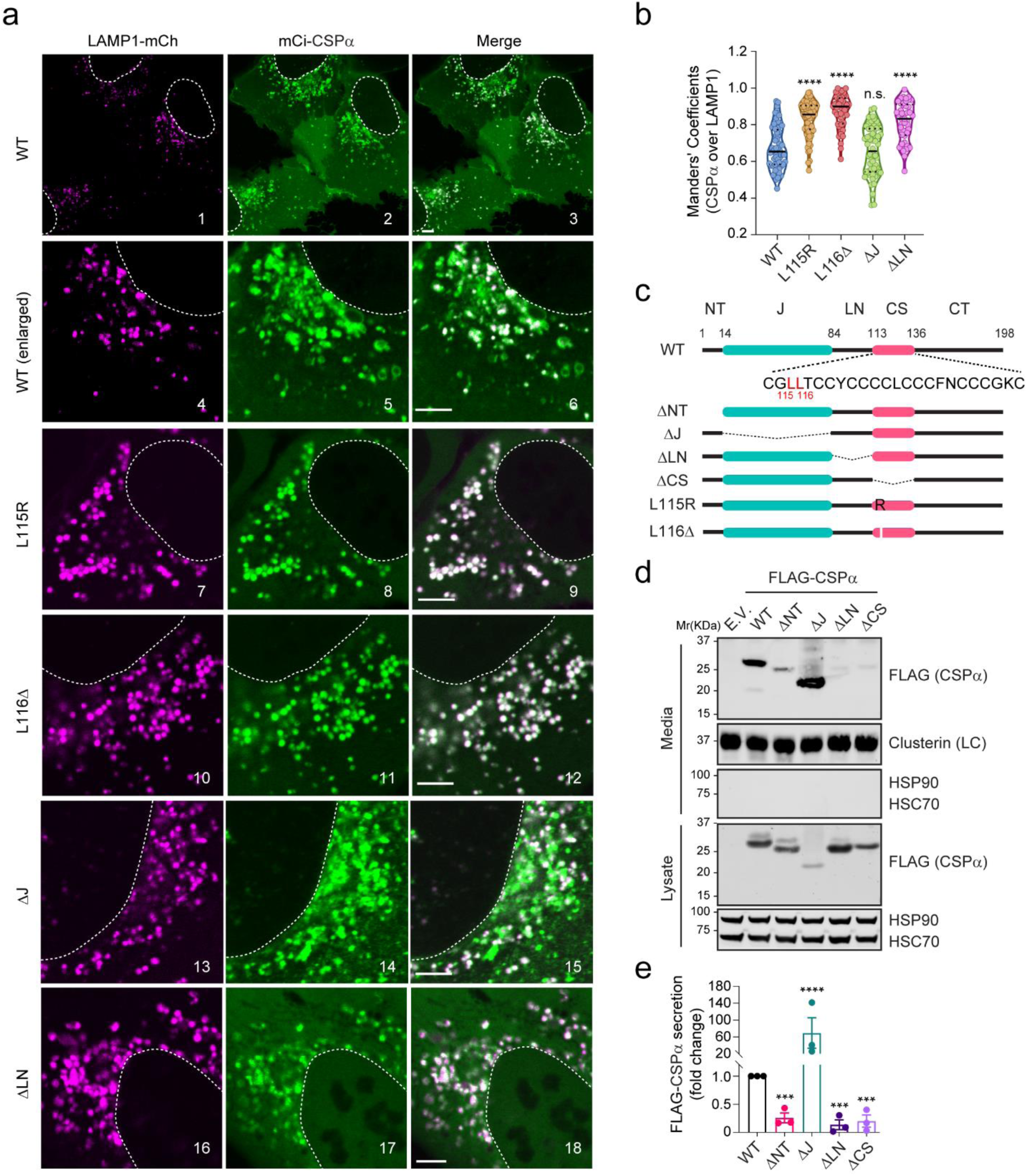
The linker domain localizes CSPα to a perinuclear LAMP1-negative compartment and is required for MAPS. **a, b,** CSPα localization to a LAMP1-negative perinuclear compartment requires the LN domain. a, Representative confocal images of U2OS cells transfected with LAMP1-mCh together with the indicated CSPα variants. Scale bars, 5 μm. b, Quantification of the co-localization efficiency of various CSPα variants with LAMP1-mCh in individual cells in a. ****, *p* < 0.0001 by one-way ANOVA plus Dunnett’s test; n.s, not significant; *n* = 60, 66, 47, 60 and 45 cells, respectively. **c,** A scheme illustrates the various CSPα variants tested in a and d. **d,** The LN domain is required for CSPα secretion. Conditioned medium and cell lysates from HEK293T cells expressing the indicated CSPα variants were analyzed by immunoblotting. LC, loading control. **e,** Quantification of the CSPα secretion experiments shown in d. Error bars indicate mean ± s.e.m. from *n*=3 independent experiments. ***, *p* < 0.0005; ****, *p* < 0.0001 (*p* for ΔNT = 0.0006, *p* for ΔLN = 0.0002, *p* for ΔCS = 0.0003) by one-way ANOVA plus Dunnett’s test.

To determine the domain(s) responsible for targeting CSPα to the LAMP1 negative vesicles, we characterized the localization of a set of CSPα deletion mutants (Fig. 4c). As previously reported ^11^, a Ci-CSPα mutant lacking the CS domain (ΔCS) was entirely localized to the cytosol due to defects in both membrane binding and palmitoylation (Supplementary Fig. 2d, e). By contrast, Ci-CSPα ΔNT (Supplementary Fig. 2d, e) and ΔJ (Fig. 4a panels 13-15, Fig. 4b) were localized to both LAMP1-positive and LAMP1-negative vesicles similarly to WT CSPα, suggesting that HSC70-binding is dispensable for this localization. Intriguingly, a Ci-CSPα mutant lacking the linker (ΔLN) was almost exclusively localized to LAMP1-positive vesicles (Fig. 4a, panels 16-18, Fig. 4b). Similar to the ANCL mutants, Keima-CSPα ΔLN also showed increased endolysosomal translocation (Fig. 3f). The similarity between the ΔLN and ANCL mutants prompted us to test the secretion of CSPα ΔLN by immunoblotting, which showed markedly reduced secretion similarly to CSPα ΔCS (Fig. 4d, e). Because CSPα ΔNT also showed reduced secretion similarly as CSPα ΔLN (Fig. 4d, e), these results suggest that the LN and CS domain act together to confer the perinuclear CSPα localization, which is necessary but not sufficient for MAPS.

### CSPα interacts with CD98hc via the linker domain

To identify the factor(s) that regulate the association of CSPα with the LAMP1-negative vesicles, we combined crosslinking with tandem affinity purification using cells stably expressing CSPα ΔJ bearing FLAG- and SBP (Streptavidin binding protein) tags (Supplementary Fig. 3a, b). We used CSPα ΔJ to avoid Hsc70 and its associated proteins. Mass spectrometry analyses of proteins co-purified with CSPα ΔJ-SBP-FLAG identified CD98hc (also named SLC3A2) as a CSPα binding partner (Fig. 5a, Supplementary Fig. 3c). CD98hc is a cofactor of heterodimeric amino acid transporters ^48^. Immunoblotting confirmed that endogenous CD98hc interacts with not only CSPα ΔJ but also WT CSPα, although the latter appeared to have a lower affinity to CD98hc than CSPα ΔJ (Fig. 5b). Reciprocal pulldown using a cell line bearing a GFP tag at the endogenous CD98hc locus further confirmed the interaction (Fig. 5c). Importantly, when we transfected various CSPα variants into *CD98hc::GFP* cells and performed the GFP pulldown, only CSPα ΔLN showed significantly reduced binding to CD98hc compared to other variants (Fig. 5c), demonstrating a specific interaction mediated by the linker domain.

**Figure 5.**
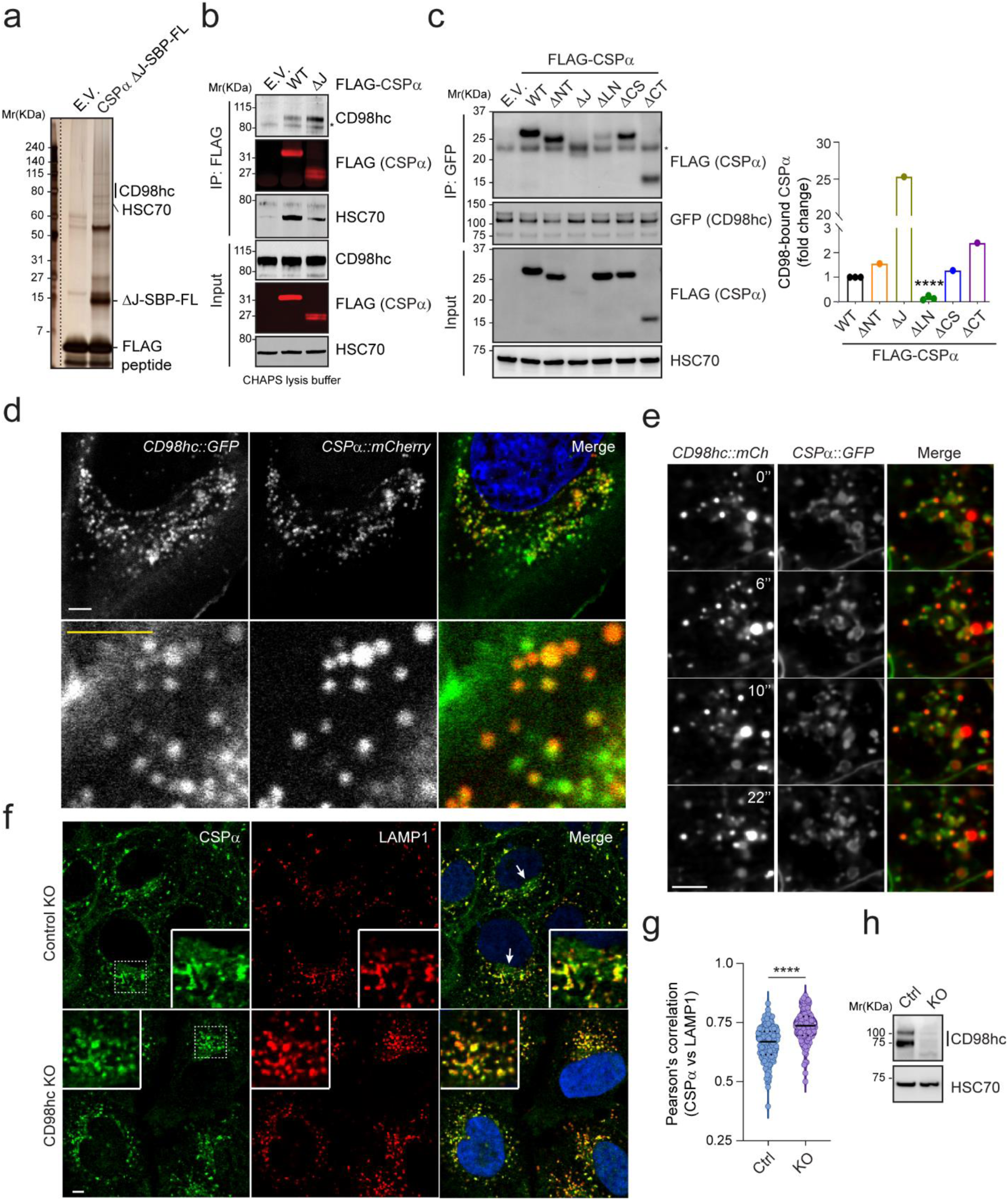
CD98hc interacts with CSPα via the LN domain and is required for the perinuclear localization of CSPα. **a,** A silver-stained gel showing the proteins co-purified with CSPα ΔJ-SBP-FLAG. **b,** Co-immunoprecipitation and immunoblotting confirm the interaction of both WT CSPα and CSPα ΔJ with endogenous CD98hc. **c,** The LN domain is required for CSPα interaction with CD98hc. *CD98hc::GFP* cells were transfected with the indicated CSPα variants for GFP pulldown and immunoblotting. Asterisk indicates IgG band. The graph shows the quantification of CD98hc-associated CSPα normalized to CSPα in cell lysates. Error bars indicate mean ± s.e.m.; ****, *p* < 0.0001 by unpaired Student’s t-test; *n*=3 independent experiments for WT and ΔLN. **d, e,** Co-localization of endogenous CSPα and CD98hc. U2OS cells (in d) or HEK293 cells (in e) bearing GFP and mCherry tag on CD98hc and CSPα respectively were imaged by dual-color confocal microscopy. e, Representative frames from Supplementary video 1. Scale bars, 5 μm. **f-h,** Depletion of CD98hc abolishes a perinuclear pool of CSPα. f, Representative images from control or CD98hc knockout (KO) U2OS cells stained by CSPα and LAMP1 antibodies. Scale bars, 5 μm. g, Quantification of the co-localization efficiency of CSPα and LAMP1 in f. ****, *p* < 0.0001 by one-way ANOVA plus Dunnett’s test; *n* = 156 and 106 cells, respectively. h, Immunoblotting confirms the depletion of CD98hc in CRISPR KO cells.

To further confirm the interaction between CD98hc and CSPα, we examined the localization of *CD98hc::GFP* by confocal microscopy. In addition to the plasma membrane, CD98hc-GFP was detected mainly on intracellular vesicles clustered around the nucleus (Supplementary Fig. 4a), a pattern similar to that of endogenous CSPα. Like CSPα, CD98hc-GFP was localized to LAMP1- and Lysotracker-positive vesicles, but many CD98hc-positive vesicles around the nucleus were free of LAMP1 (Supplementary Fig. 4b-b’’). As anticipated, cells expressing GFP and mCherry on endogenous CSPα and CD98hc respectively showed extensive co-localization of the two proteins on vesicles over time (Fig. 5d, e, Supplementary video 1), which supports the identified interaction.

### CD98hc is required for the perinuclear association of CSPα and MAPS

The fact that CSPα ΔLN is defective in CD98hc interaction and absent from the LAMP1-negative perinuclear compartment suggests CD98hc as a potential regulator of the CSPα localization. To test this idea, we knocked out CD98hc in U2OS and HEK293T cells and immunostained endogenous CSPα and LAMP1 in CD98hc-depleted and control knockout cells. While in control cells, the presence of CSPα in the LAMP1 negative compartment was readily visible, in CD98hc KO cells, CSPα was almost completely absent from this compartment (Fig. 5f-h; Supplementary Fig. 5a, b), while its co-localization with LAMP1 became more prominent. By contrast, the plasma membrane-associated CSPα was not affected in CD98hc-depleted cells (Supplementary Fig. 5a, b). Flow cytometry showed that CD98hc deficiency or overexpression of CD98hc did not significantly affect the endolysosome translocation of Keima-CSPα (Supplementary Fig. 5c, d). Thus, CD98hc is required for the perinuclear association of CSPα but dispensable for CSPα endolysosome translocation and plasma membrane association.

We next tested whether CD98hc is required for CSPα-mediated MAPS using α-Syn and GFP1-10 as model substrates. Indeed, the secretion of these proteins under basal conditions or in CSPα-overexpressing cells was significantly reduced when CD98hc was depleted (Fig. 6a-d; Supplementary Fig. 5e-g). Moreover, the secretion of CSPα itself was also dramatically inhibited in CD98hc deficient cells (Fig. 6a, b, e). These results strongly suggest that CD98hc is required for CSPα-mediated MAPS.

**Figure 6.**
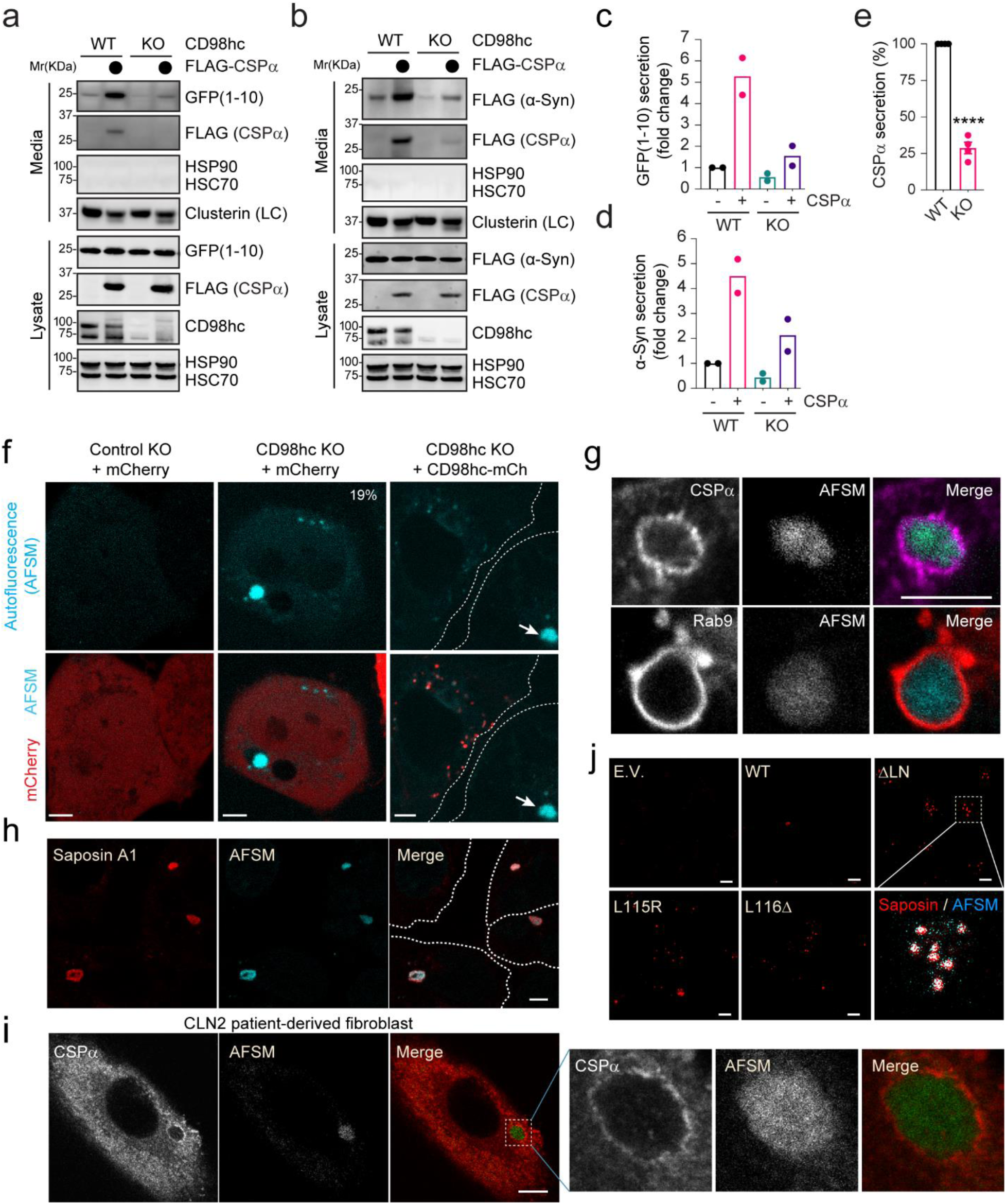
Depletion of CD98hc inhibits MAPS and causes the accumulation of AFSMs in cells. **a,** Depletion of CD98hc inhibits GFP1-10 secretion. Conditioned medium and cell lysates from the control or CD98hc KO cells transfected with GFP1-10 and the indicated plasmids were analyzed by immunoblotting. **b,** As in a, except that cells transfected with FLAG-α-Syn and the indicated plasmids were analyzed. **c,** Quantification of the experiments in a (*n* = 2). **d,** Quantification of the experiments in b (*n* = 2). **e,** Quantification of the CSPα secretion from the experiments in a and b. Error bars indicate mean ± s.e.m.; ****, *p* < 0.0001 by unpaired Student’s t-test; *n* = 4 independent experiments. **f-i,** CD98hc deficient cells contain AFSMs resembling NCL-associated lipofuscin. f, Control (Ctrl.) KO or CD98hc KO HEK293T cells were transfected with mCherry (mCh) or CD98hc-mCh and imaged by confocal microscopy. The dashed lines indicate cell boundary. Arrow indicates a “lysobody” in a cell without CD98hc-mCh. g, Enlarged images of CD98hc KO cells stained by either CSPα antibodies (Top panels) or transfected with mCh-Rab9 (Bottom panels) show that AFSMs are surrounded by CSPα- and Rab9-positive membranes. h, CD98hc KO U2OS cells stained with Saposin A1 antibodies in red. i, Fibroblast cells from NCL2 patients were stained with CSPα antibodies. j, HEK293T cells transfected with the indicated CSPα variants were stained by Saposin A1 antibodies in red. Scale bars, 5 μm.

The link of perinuclear CSPα to unconventional secretion prompted us to investigate whether CSPα is co-localized with TMED10, a recently identified unconventional protein secretion regulator that is localized to ERGIC vesicles ^49^. We endogenously tagged CSPα with GFP and TMED10 with mCherry. Dual-color fluorescence microscopy showed that the perinuclear CSPα is not co-localized with TMED10 in U2OS cells (Supplementary Fig. 2c).

### Depletion of CD98hc causes lipofuscin-like structures in mammalian cells

Given the link between *CSPα* mutations and lipofuscinosis, we sought to determine whether CD98hc deficiency also caused the accumulation of lipofuscin-like autofluorescent storage materials (AFSM). Indeed, confocal microscopy showed that ∼20% of CD98hc KO cells contained one or two large spherical punctae detectable by excitation with a UV light (Fig. 6f, Supplementary Fig. 6a). By contrast, AFSMs were only detected in less than 0.5% of WT cells. Importantly, this structure was not present when CD98hc was re-expressed in the KO cells, indicating that the lack of CD98hc leads to AFSM formation (Fig. 6f). 3D confocal analysis combined with dual-color fluorescence microscopy showed that these AFSMs were in enclaves surrounded by membranes enriched for CSPα and Rab9, suggesting their origin from endolysosomes (Fig. 6g; Supplementary Fig. 6b). Additional immunostaining detected Saposin A1, a major known component of lipofuscin ^50^ in all AFSMs observed in CD98hc KO cells (Fig. 6h), confirming their identity as lipofuscin. Additionally, cells derived from late INCL patients bearing *Cln2* mutations also contained large lipofuscin-like AFSMs morphologically identical to those in CD98hc KO cells^51^ and were also wrapped around by CSPα-positive membranes (Fig. 6i). Interestingly, cells overexpressing CSPα L115R or L116Δ also contained an increased number of Saposin A1-positive AFSMs, although they were often smaller in size compared to those in CD98hc KO cells (Fig. 6j). Because *Cln2* encodes a lysosomal peptidase TPP1, these observations suggest that lipofuscin biogenesis is associated with either defective lysosomal digestion or excessive accumulation of proteins and membranes into endolysosomes (see discussion).

### AFSM accumulation and CD98hc deficiency contribute to neurodegeneration in a fly ANCL model

We next used a recently established fly model to further evaluate the role of CSPα-associated AFSM and CD98hc in neurodegeneration. We expressed WT human CSPα or the ANCL-associated L116Δ mutant in the developing *Drosophila* larval eyes using the GMR-Gal4 driver. As reported previously ^38^, CSPα L116Δ overexpression caused massive neuronal cell death, resulting in a severe rough eye phenotype in adult flies (Fig. 7a’). By contrast, WT CSPα did not change the eye morphology at 25 °C (Fig. 7a). Interestingly, confocal microscopy analyses of larval eye discs showed that the CSPα L116Δ-expressing tissues contained many autofluorescent punctae in the apical cytoplasm (Fig. 7b, b’ and c, c’) within the photoreceptor cells (Fig. 7d).

**Figure 7.**
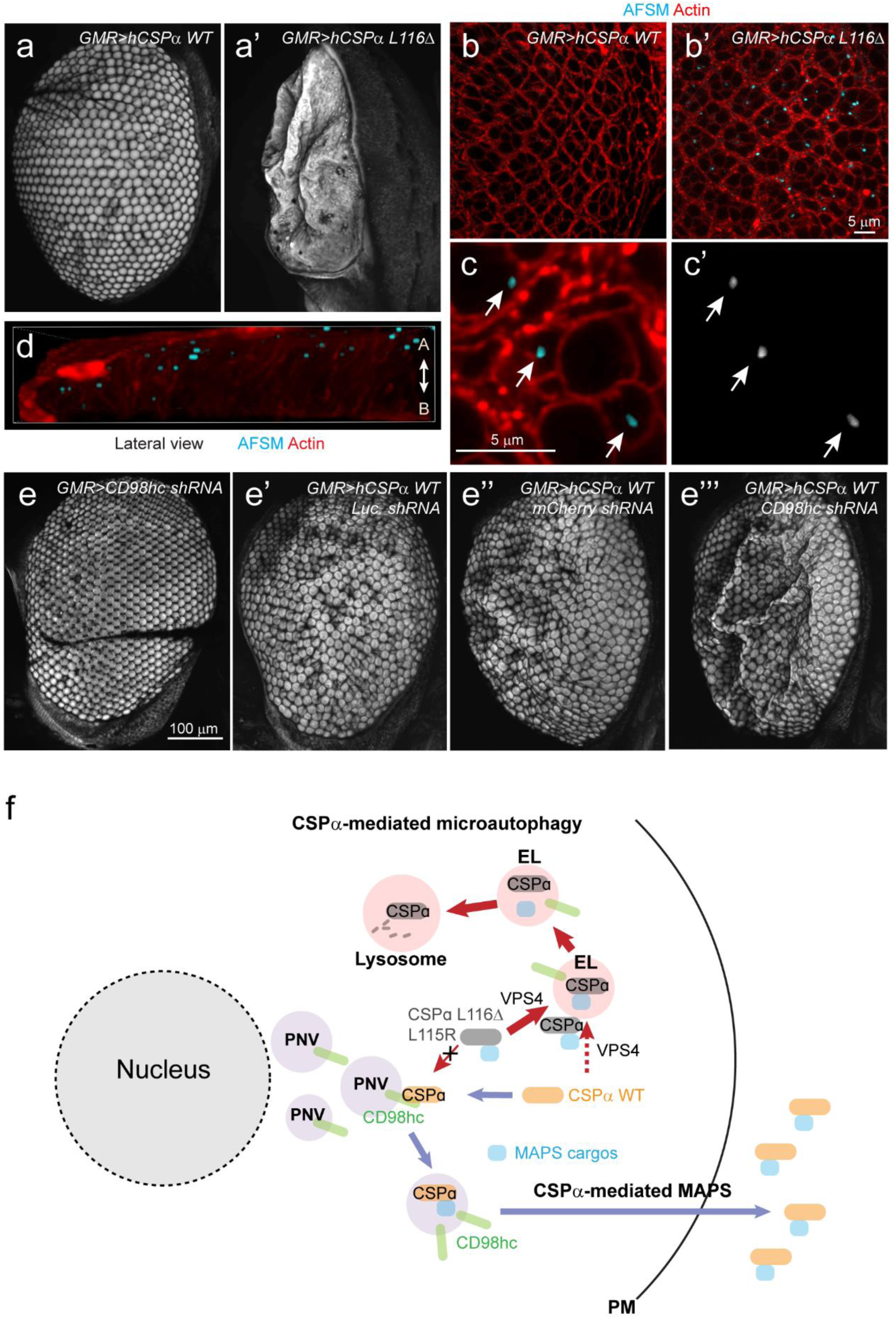
AFSM accumulation and genetic interaction between CSPα and CD98hc in a fly ANCL model. **a-a’,** The expression of human CSPαL116Δ but not CSPα WT in fly photoreceptor cells causes a severe rough eye phenotype. **b-d,** AFSM accumulation in photoreceptor cells in larvae expressing human CSPα L116Δ. b-b’, Imaginal eye discs from GMR>hCSPα WT or GMR>hCSPα L116Δ were stained with phalloidin to label cortex actin (red) and imaged at Ex_405_ to detect AFSM. c, c’, An enlarged view of a cluster of photoreceptor cells bearing AFSMs (arrows) in the cytoplasm. d, A 3D side view of AFSM in a CSPα L116Δ-expressing eye disc. A, apical; B, Basal. **e-e’’’** CD98hc knockdown enhances the rough eye phenotype associated with hCSPα WT expression at 28 °C. **f,** The CSPα-mediated protein triaging pathways, and the deregulation of these processes in cells expressing ANCL-associated CSPα mutants. PM, plasma membrane; EL, endolysosome; PNV, perinuclear vesicles.

Because CD98hc regulates the CSPα function in MAPS, if disruption of MAPS while maintaining CSPα-mediated microautophagy contributes to AFSM production and neurodegeneration, knockdown of CD98hc should enhance CSPα-induced neurodegeneration. We, therefore, crossed *GMR>hCSPα* WT and *GMR>hCSPα* L116Δ flies to a strain bearing a CD98hc-targeting shRNA downstream of the UAS regulatory element, or as controls to strains bearing either a luciferase shRNA or mCherry shRNA. When raised at 28 °C, flies expressing CD98hc shRNA alone had largely normal eyes (Fig. 7e), whereas moderate rough eye phenotype was observed in flies expressing WT hCSPα together with a control shRNA (Fig. 7e’, e’’). By contrast, flies expressing WT CSPα together with CD98hc shRNA had more severe rough eyes (Fig. 7e’’’) with reduced eye size and a significant loss of the pigments (Supplementary Fig. 7b-b’’). Likewise, knockdown of CD98hc also enhanced the rough eye phenotype in hCSPα L116Δ- expressing flies cultured at 25 °C (Supplementary Fig. 7c-c’’). Collectively, these results support a model that loss of MAPS activity while maintaining CSPα-mediated microautophagy disrupts lysosome homeostasis, causing AFSM accumulation and neurodegeneration (Figure 7f).

## Discussion

In this study, we show that CSPα couples two protein quality control processes to limit microautophagy, which is essential for endolysosome homeostasis. *CSPα* mutants uncouple these processes, causing lipofuscinosis and neurodegeneration. Specifically, we show that a fraction of CSPα is translocated into endolysosomes together with its clients. This process is analogous to previously reported selective microautophagy (also known as endosomal microautophagy or eMI), which is mediated by HSC70 and ESCRT proteins ^44, 45, 47^. Like eMI, CSPα-mediated microautophagy also involves the ESCRT machinery, but it is not dependent on Hsc70. Given that CSPα cargos are misfolded proteins, other CSPα-associated chaperones such as SGT ^16^ may assist it in substrate recruitment. Intriguingly, recent studies showed that the L115R and L116Δ mutations reduce CSPα palmitoylation ^39^. These mutants are also more prone to aggregation ^37, 38, 52^. Nevertheless, these mutants have increased endolysosome translocation activities compared to WT CSPα, suggesting that CSPα-mediated microautophagy does not require palmitoylation. Indeed, several CSPα palmitoylation defective mutants can still bind to membranes in cells ^11^.

A fraction of CSPα is also localized to a perinuclear membrane compartment, which is largely free of LAMP1 and not stained well by a Lysotracker dye. However, this compartment can be weakly labeled by Lysotracker after prolonged staining (Supplementary video 2), suggesting that it might be a pre-lysosomal compartment less acidic than the lysosomes. The localization of CSPα to this compartment is regulated by the newly identified CSPα binding partner CD98hc. CD98hc is a type II membrane glycoprotein capable of interacting with six light chain molecules, forming a set of amino acid transporters on the plasma membrane ^48^. Moreover, additional adaptors such as LAPTM4b can retain a CD98hc-containing transporter in endolysosomes, regulating the amino acid balance between the endolysosomes and cytoplasm and thus the activation of mTOR signaling ^53^. Whether CD98hc-dependent regulation of the CSPα perinuclear localization requires additional components or a complete amino acid transporting system awaits further characterization.

Although the CSPα-associated pre-lysosomal compartment does not overlap with TMED10, an ER-Golgi intermediate compartment (ERGIC) protein recently implicated in unconventional protein secretion, several lines of evidence suggest that this compartment functions in MAPS. First, the CSPα mutants defective in binding to this compartment all fail to promote MAPS. Additionally, the knockout of CD98hc impairs the association of CSPα to this compartment, which also reduces the secretion of misfolded proteins. In *S. Cerevisiae*, unconventional protein secretion under stress conditions is mediated by a Golgi-derived compartment termed CUPS ^54, 55^. The pre-lysosomal CSPα-positive compartment may be functionally analogous to CUPS. In yeast, protein translocation into CUPS is thought to be mediated by the ESCRT machinery and some autophagy regulators ^55^. By contrast, unconventional secretion of misfolded proteins in mammalian cells is not dependent on ESCRT and cannot be blocked by the autophagy inhibitor 3-MA ^32^. How misfolded proteins enter the CSPα-associated perinuclear compartment remains to be elucidated.

CSPα-mediated microautophagy and MAPS appear to operate in parallel to process misfolded proteins. Conceivably, these processes, when properly tuned, should reduce misfolded proteins to improve protein homeostasis. By contrast, deregulation of these processes may lead to an overflow of misfolded proteins to either endolysosomes or cell exterior. The fact that CSPα lacking the J-domain has significantly increased activities in both endolysosomal translocation and protein secretion suggests an autoinhibitory mechanism that tightly controls these processes. Consistently, structural studies have illustrated phosphorylation-dependent conformational changes, which disrupt a J-domain-mediated intermolecular interaction ^56^.

Our findings suggest that ANCL-associated CSPα mutations cannot be classified as simple loss- or gain-of-function alleles. Instead, while these mutations abolish the CSPα pre-lysosomal localization and the corresponding MAPS function, they increase the translocation of CSPα into lysosomes. One presumed consequence is the abnormal flow of misfolded proteins and membranes into lysosomes, damaging this organelle over time and resulting in undigested remnants in the form of lipofuscin. This model is consistent with the finding that mutations in PPT1 (palmitoyl protein thioesterase 1) also result in similar disease phenotypes ^57^. Presumably, endolysosome-associated CSPα is processed by PPT1, which may regulate its function in proteostasis regulation. While our study suggests that inappropriate triaging of misfolded proteins by the two CSPα-mediated protein trafficking branches can generate lipofuscin and cause neurodegeneration, additional studies are needed to elucidate how these processes are precisely controlled under physiological conditions to avoid proteostasis catastrophes.

## Methods

### Cell lines, siRNAs, Plasmids, and Antibodies

HEK293T, HEK293, COS-7, and U2OS cells were purchased from ATCC. Human patient-derived CLN2 fibroblast cell is a gift from Wei Zhang (NCATS/NIH). The cells were maintained in DMEM medium (Corning cellgro) supplemented with 10% fetal bovine serum (Gibco) and penicillin-streptomycin (Gibco). Transfection was performed with TransIT-293 (Mirus) for HEK293T cells, and with Lipofectamine2000 (Invitrogen) for COS-7 and U2OS cells. Lipofectamine RNAiMAX (Invitrogen) was used in all gene silencing experiments according to the manufacturer’s protocol. HEK293T and U2OS cells were used to stably express or were depleted of target proteins by lentivirus infection and selection for 3-7 days, using puromycin (0.3 μg/mL for HEK293T, 1 μg/mL for U2OS) or hygromycin (200 μg/mL). To generate CD98hc CRISPR knockout HEK293T cell lines, we constructed two sgRNA-CAS9 guide constructs in pX330 hCas9 D10A based on a published protocol ^58^. The primers containing targeting sequences for the human CD98hc gene are:

Target 1, forward primer: 5’-caccGCTGCAGATCGACCCCAATTT-3’;
Target 1, reverse primer: 5’-aaacAAATTGGGGTCGATCTGCAGc-3’;
Target 2, forward primer: 5’-caccGTCTGAGCGACATCATCCTTC-3’;
Target 2, reverse primer: 5’-aaacGAAGGATGATGTCGCTCAGAc-3’

The two CD98hc targeting constructs were co-transfected into HEK293T cells. At 24 h post-transfection, cells were diluted and seeded into a 96-well plate at <1 cell per well. Single cell-derived clones were screened for CD98hc deficiency by immunoblotting. To generate CD98hc CRISPR knockout U2OS cell line, a sgRNA guide sequence (the Target 1 sequence above) was cloned into pLenti CRISPRv2 vector, and the lentivirus was used for infection as described previously ^59^.

Endogenous tagging was performed as described previously ^60^ (http://www.pcr-tagging.com). Briefly, the PCR cassettes were amplified from pMaCTag-05 plasmid by the AccuPrime™ Pfx DNA polymerase using the primers listed below:

M1_CSPα, forward primer: 5’- ACGCCGATCGTCATACAGCCGGCATCCGCCACCGAGACCACCCAGCTCACAGCCGACTCCCACCCCAGCTACCACACTGACGGGTTCAACTCAGGTGGAGGAGGTAGTG-3’
M2_CSPα_enAsCas12a, reverse primer: 5’- TAAGGTTGGCGTGGCCAGGCGCCGGCTCCTCCTCTGACCACAGCTCCTCCTGGATAAAAAAAGCTCCTCCTGGATTTAGTTATCTACAAGAGTAGAAATTAGCTAGCTGCATCGGTACC-3’
M1_CD98hc, forward primer: 5’- CCAGGCCGTGAGGAGGGCTCCCCTCTTGAGCTGGAACGCCTGAAACTGGAGCCTCACGAAGGGCTGCTGCTCCGCTTCCCCTACGCGGCCTCAGGTGGAGGAGGTAGTG-3’
M2_CD98hc_enAsCas12a, reverse primer: 5’- GCCAAAGGGCCTGGGAAGGAAAGGAGAAGGGTAGTGGGTCCATGTCAGGCTGAAGAAAA AATGAAGTCAGGCCGCGTAGGGATCTACAAGAGTAGAAATTAGCTAGCTGCATCGGTACC-3’
M1_TMED10, forward primer: 5’- AGCATCTTTTCAATGTTCTGTCTCATTGGACTAGCTACCTGGCAGGTCTTCTACCTGCGACGCTTCTTCAAGGCCAAGAAATTGATTGAGTCAGGTGGAGGAGGTAGTG-3’
M2_TMED10_enAsCas12a, reverse primer: 5’- CCAGCGATGTTCTGCTGGCTGAGGTACAAGGTGGGAGGAGAATATGCCTCATTCAAAAAAATGCCTCATTCATTACTCAATATCTACAAGAGTAGAAATTAGCTAGCTGCATCGGTACC-3’

The PCR products were gel purified using a QIAGEN Gel Extraction Kit. HEK293T cells were transiently transfected with 1 µg of a PCR cassette and 1 µg of pCAG- enAsCas12a(E174R/S542R/K548R)-NLS(nuc)-3xHA (gift from Keith Joung & Benjamin Kleinstiver, Addgene plasmid # 107941) using TransIT293 (Mirus) according to the manufacturer’s protocol and GFP or mCherry positive cells were sorted two weeks later by FACS. For making double tagging cells (*CSPα::GFP / CD98hc::mCherry*), The PCR cassette for CD98hc described above was transfected to *CSPα::GFP* cell and cells showing double fluorescences were sorted by FACS. All expression constructs, si-RNAs, chemicals, and antibodies are listed in Supplementary Table 1.

### Protein samples preparation and Immunoblotting

Conditioned media and cell lysates were prepared as described previously ^60^. Immunoblottings were performed using the standard protocols. All primary antibodies were diluted in 5% BSA in phosphate buffer saline (PBS) as described in Supplementary Table 1. To quantify secreted protein in conditioned media, HRP-conjμated secondary antibodies were used. Immunoblotting signal was detected by the enhanced chemiluminescence method (ECL) and recorded by a Fuji LAS-4000 imager or Bio-Rad Chemidoc. The intensity of the detected protein bands was quantified by ImageGauge v3.0, Bio-Rad ImageLab, or ImageJ. Protein secretion levels were determined by normalizing the level of the secreted proteins by the amount of the same protein in cell lysates. For other immunoblotting quantifications, fluorescently labeled secondary antibodies were used. Immunoblots were scanned by a LI-COR Odyssey scanner or Bio-Rad Chemidoc.

### Membrane fractionation, Proteinase K protection assay, and Immunoprecipitation

To fractionate cells, five million mKeima-CSPα HEK293T cells were harvested, washed with ice-cold PBS, and treated with 300 μl PB buffer (115 mM KOAc, 5 mM NaOAC, 25 mM HEPES pH 7.3, 2.5 mM MgCl_2_, 0.5 mM EGTA) containing 0.05% digitonin and 1 mM DTT plus a protease inhibitor cocktail for 5 min on ice. The semi-permeabilized cells were confirmed by Trypan blue staining before proceeding with the following steps. Cytosol fractions were saved from the supernatant after cells were centrifuged at 17,000 x g for 5 min at 4 °C. The remaining membrane pellets were washed with 1x PB buffer without digitonin once and resuspended in 300 μl of 1x PB buffer by gentle pipetting. Twenty microliters of the membrane fractions were incubated with 5 μl of proteinase K (Thermo, 20 mg/mL) pre-diluted at either 1:750 or 1:1,500 for 12 min at room temperature. Reactions were quenched by adding a preheated 4x Laemmli buffer and boiling for 20 min. The samples were analyzed by SDS-PAGE electrophoresis followed by immunoblotting. For co-immunoprecipitation assays, pre-equilibrated FLAG M2 agarose beads (Sigma) or GFP-trap beads (Chromotek) were incubated with lysates containing tagged proteins for 1 hour at 4 °C. The beads were washed with the lysis buffer, and the bound proteins were eluted in 1x Laemmli buffer at 95 °C and resolved by SDS-PAGE for immunoblotting analysis.

### Tandem affinity purification and mass spectrometry of FLAG-SBP-CSPα precipitates

A schematic flow chart of the purification procedure is presented in Supplementary Figure 3. Three p150 dishes of HEK293T cells, which were transfected with 15 μg of either empty vector, FLAG-Streptavidin binding peptide (SBP)-tagged CSPα WT or CSPα ΔJ, were harvested in PBS. After centrifugation at 1,000 x g for 10 min at 4 °C, the cell pellets were treated with 0.025 % digitonin in the PB buffer containing 1 mM DTT and a protease inhibitor cocktail for 5 min. Once membrane permeabilization is confirmed by trypan blue staining, the lysates were spun at 16,000 g for 5 min. The resulting membrane pellets were washed with 1x PB buffer followed by centrifugation. The washed pellets were resuspended in 0.33 % formaldehyde in PB buffer and incubated at 37 °C for 25 min with rocking for crosslinking. After top-speed centrifugation for 5 min, the pellets were further lysed in 8 mL of RIPA lysis buffer with 1 mM DTT and a protease inhibitor cocktail for 1 h at 4 °C. After centrifugation for 5 min, the supernatants were collected for the following tandem affinity purification steps.

To purify CSPα and its interacting proteins, the RIPA-soluble fractions were incubated with 200 μl of pre-washed FLAG M2 agarose bead slurry with rocking for 1 h at 4 °C. The beads were washed with 1 mL of RIPA buffer three times and 1 mL of streptavidin binding buffer (10 mM Tris-HCl pH7.5, 1 mM EDTA, 1M NaCl) once. After spinning down, the bound proteins were eluted by incubation with 250 μl of 3 x FLAG peptide solution (200 μg/mL) (Sigma) for 15-20 min twice. After centrifugation, the elutes (∼500 μl) were incubated with an equal volume of streptavidin binding buffer and 80 μl of pre-washed streptavidin beads (Thermo). After incubation for 1hr at 4 °C, the beads were washed with streptavidin binding buffer three times. The bound proteins were eluted in 40 μl 1x Laemmli buffer by boiling for 20 min. The purified proteins were resolved by SDS-PAGE and visualized by silver staining using the SilverQuest kit (Invitrogen) according to the manufacturer’s protocol. Mass spectrometry analysis of CSPα co-purified proteins in gel slices was done by the Taplin Mass Spectrometry facility of Harvard Medical School by a fee-based service.

### Imaging analysis

For immunostaining, cells cultured on 1.5 thickness coverslips precoated with poly-D-lysine (50 μg/mL) were fixed in 4% paraformaldehyde in PBS or in cold methanol for 10 min. Cells were then washed with PBS twice and permeabilized with a PBS-based staining solution containing 0.2% saponin and 10% FBS for 10 min at room temperature. Cells were stained by primary antibodies diluted in the staining solution overnight at 4 °C and washed with PBS three times. Alexa Fluor® secondary antibodies (Thermo) were diluted in the staining solution and added to cells for 1 h at room temperature. If necessary, DAPI (Sigma, 1 μg/mL) was included in the staining solution to label nuclei. After washing with PBS three times, the coverslips were mounted on glass slides with mounting media (Vectashield). For live-cell imaging, cells were seeded to μ-slide chamber (ibidi) precoated with fibronectin (10 μg/mL, Sigma) for 1 hr. Culture media were replaced with imaging medium (phenol-red free DMEM containing 10% FBS) before imaging. Cells were stained with Lysotracker red (Thermo) or Alexa^594^-labeled cholera toxin subunit B (CTB, Biotium) according to manufacturer’s protocols. AFSM images were detected using a DAPI filter (Ex405/Em427-487). Photobleaching experiments were performed as described previously ^32^. Cell imaging experiments were conducted on either a LSM780 laser scanning confocal microscopy (Zeiss) or a SoRa spinning disk super-resolution microscopy (Nikon) equipped with heating and a CO_2_ incubation system. Images were further processed for dot number counting or fluorescence ratio determination using ImageJ. For colocalization assay, the co-localization tools in the NIS-elements software (Nikon) or JACoP plugin in ImageJ were used.

### Flow cytometry

Keima-expressing cells were dissociated into fresh DMEM medium by gentle pipetting and passed through a cell strainer cap filter (Thermo). Flow cytometry was performed on an LSRII Fortessa analyzer (Becton Dickinson). The gate for acidic (Ex586/Em620) / neutral (Ex440/Em620) intensities of individual cells (>10,000 cells) were determined manually by reference of bafilomycin A1 (100-200 nM for 2-4 h) treated samples, which converts ∼99% of cell population to the neutral gate. Flow data were analyzed using FlowJo 10.6 software (FlowJo LLC).

### Fly experiments

Fly strains bearing shRNA-expressing cassettes downstream of UAS sequences are 31603, 35785, 57746 from the Bloomington Drosophila Stock Center. The flies expressing either WT human CSPα or CSPα L116Δ were described previously ^38^. Unless otherwise specified, cultures were maintained in 25 °C incubators on standard medium. For imaging fly eyes, 5-10 adult flies were fixed in PBS containing 4% formaldehyde for 1 h, rinsed with PBS. The flies were then dehydrated by soaking sequentially in 30%, 50%, 70%, 90%, and 100% ethanol. Dried flies were mounted in an Ibidi imaging chamber and scanned by a Nikon C1 spinning disk confocal microscope using Ex. of 488 nm and Em. of 520 nm. Shown is the maximum projection view of the scanned Z-section images.

To detect AFSM in photoreceptor cells, imaginal eye discs were dissected from third instar larvae, fixed in 4% formaldehyde in PBS for 20 min at room temperature. Eye discs were washed three times with PBS, then permeabilized in a PBS-based staining solution containing 0.2% Saponin and 10% FBS for 10 min. The discs were then stained by Alexa594-labeled phalloidin (Thermo) in the staining solution for 30 min at room temperature. The discs were washed twice by PBS and then imaged by a Zeiss LSM780 laser scanning confocal microscope.

### Data analysis

All experiments were repeated two or more times. For statistical analyses, at least three independent experiments were carried out. For quantification of microscope images, at least 30 randomly selected cells were analyzed. Statistical significance was evaluated by a two-tailed t-test or one-way ANOVA followed by multiple comparison analyses of variance by Dunnett’s test using the GraphPad Prism 9 software. Differences were considered significant at the 95% level of confidence.

## Acknowledgments

We thank X. Qi (University of Cincinnati) for Saposin A1 antibodies, the Harvard Taplin Mass Spectrometry, the NHLBI flow cytometry, and the NIDDK confocal microscopy imaging cores for services, R. Puertollano (NHLBI) and J. Bonifacino (NICHD) for critical reading of the manuscript. This research was supported by an intramural research program of the NIDDK (Y. Ye) in the National Institutes of Health and by 1R21NS117855-01 to K. Zinsmaier at the University of Arizona.

## Author contributions

J.L. designed and performed most experiments and analyzed the data. Y.X. and S.L. made some key reagents. C.Z. provided fly strains and advised on the fly study. M.X. analyzed the CLN2 patient cells. Y.Y. performed the fly experiments and oversaw the study. J.L and Y.Y. wrote the paper.

## Supplementary Information

**Figure S1, related to Figure 2.**
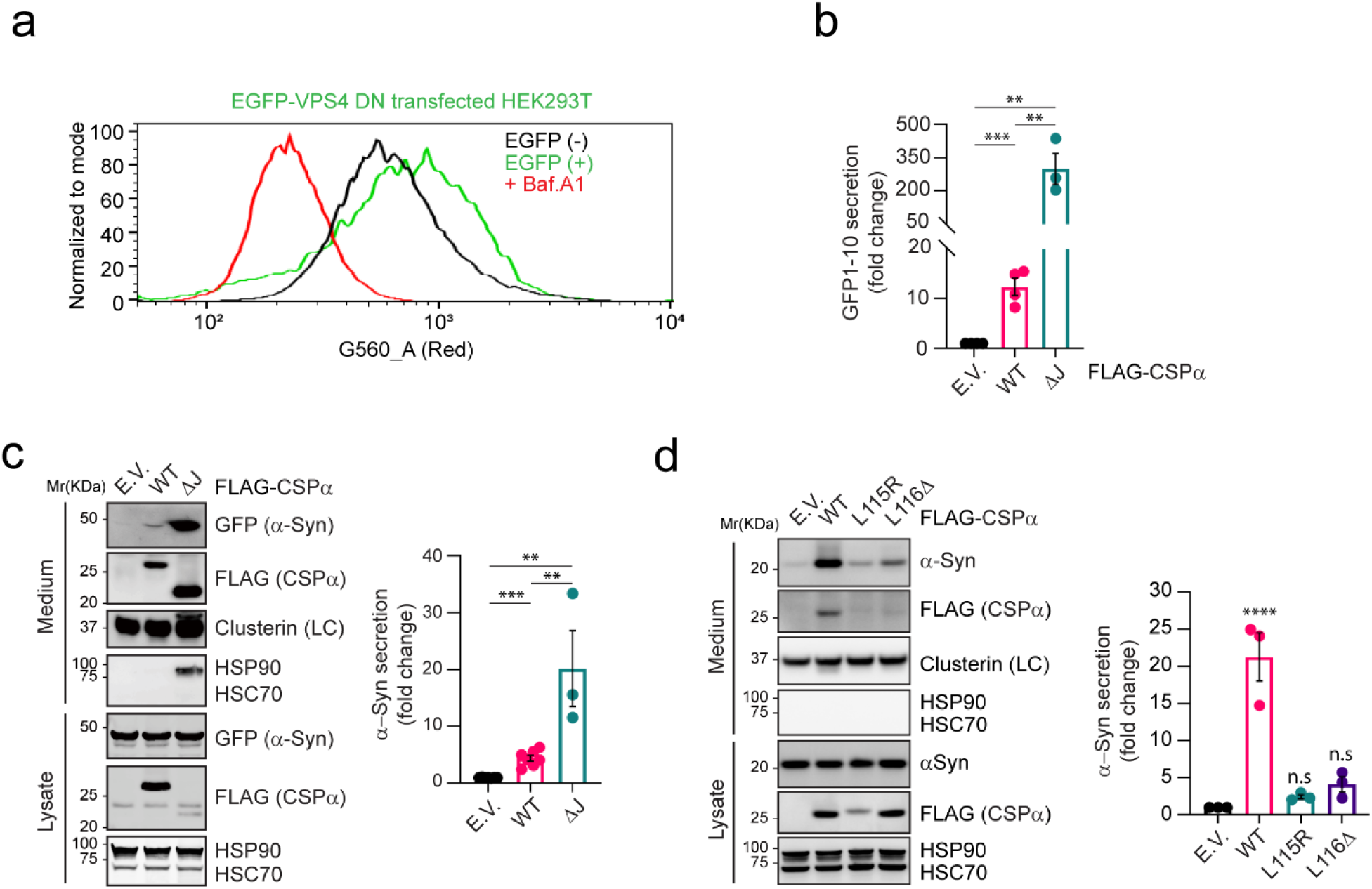
CSPα-mediated microautophagy is dispensable for MAPS. **a,** VPS4 DN expression does not increase the lysosomal pH. HEK293T cells transfected with EGFP-VPS4 DN (E228Q) were stained with a LysoTracker dye and subjected to flow cytometry analysis. Two gates were selected for comparison of EGFP negative and positive cells. Bafilomycin A1 (200 nM for 2 h) treatment was used as the positive control. **b,** Quantification of the experiments as shown in Fig. 2g; error bars, mean ± s.e.m. for fold changes in the ratio of GFP1-10 _Med_ / GFP1-10 _Lys.._ *n=* 4 experiments; *p-value* for EV/WT = 0.0006, *p-value* for EV/ΔJ = 0.0037, *p-value* for WT/ΔJ = 0.0044 by unpaired Student’s t-test. **c,** Secretion of EGFP-α-Syn is stimulated by CSPα WT and CSPα ΔJ. HEK293T cells expressing GFP-α-Syn were transfected with empty vector (EV) or CSPα plasmids. Secreted proteins were collected for 16 h and analyzed by immunoblotting as indicated. The graph shows mean ± s.e.m. for fold change in the ratio of α-Syn _Med_/ α-Syn _Lys_ from *n* =7 experiments. *p-value* for EV/WT < 0.0001, for EV/ΔJ = 0.0014, for WT/ΔJ = 0.0047 by unpaired Student’s t-test. **d,** CSPα ANCL mutants fail to induce α-Syn secretion. HEK293T cells expressing FLAG-α-Syn were transfected with the indicated CSPα plasmids. Conditioned medium (16 h) and cell lysates were analyzed by immunoblotting. The graph shows mean ± s.e.m. for fold changes in the ratio of α-Syn _Med_/ α-Syn _Lys_ from *n* = 3 experiments; ****, *p* < 0.0001; n.s. not significant by one-way ANOVA plus Dunnett’s test. LC, loading control.

**Figure S2, related to Figure 4.**
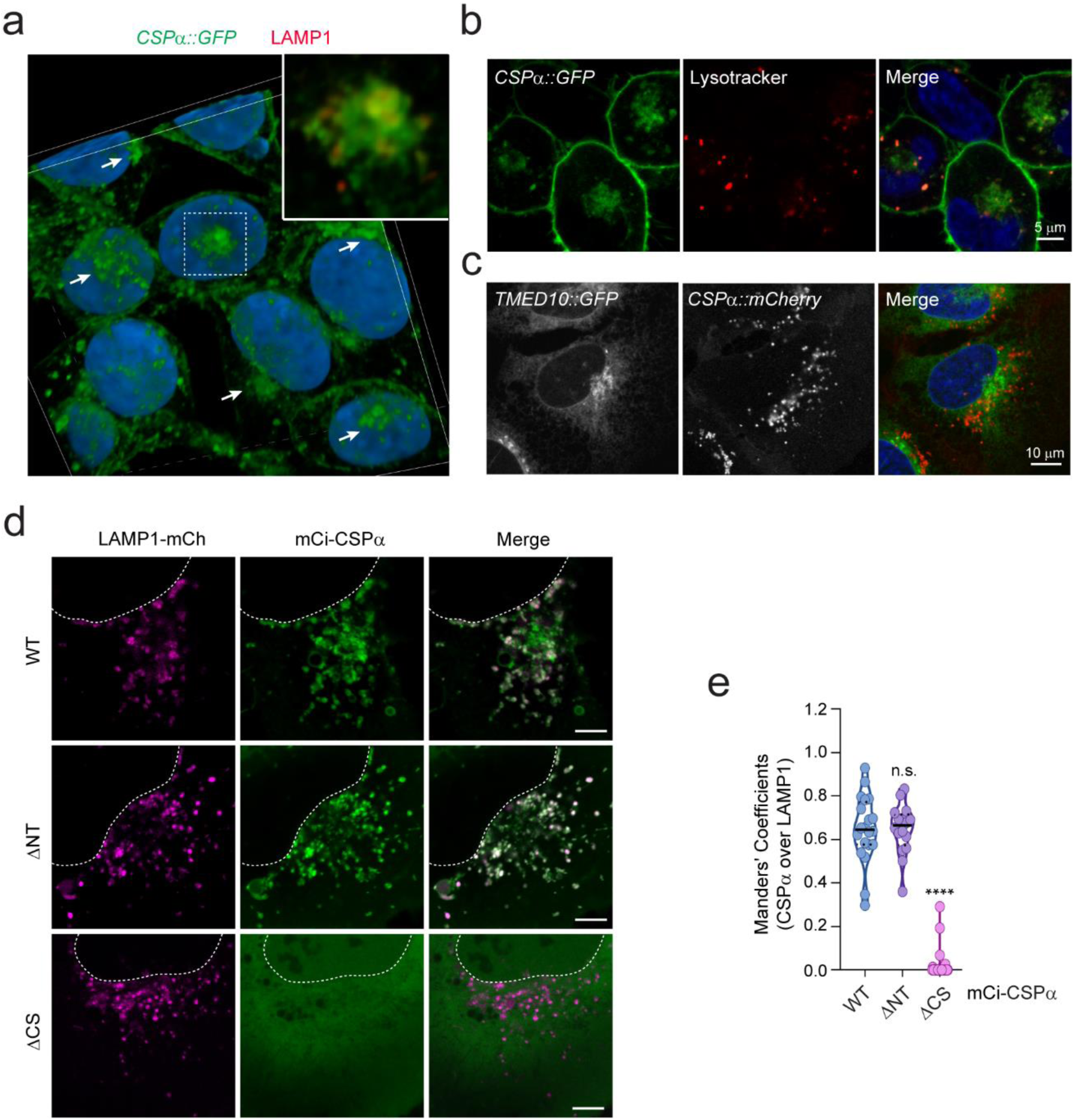
The linker domain localizes CSPα to a perinuclear LAMP1-negative compartment and is required for MAPS. **a,** A 3-D view of the subcellular localization of endogenous LAMP1 and CSPα in HEK293T cells. HEK293T cells expressing CSPα bearing an endogenously tagged GFP at the carboxyl terminus (*CSPα::GFP*) were stained with LAMP1 antibodies in red and DAPI in blue. Confocal images of z-stacks were reconstituted into a 3D view using the Nikon element software. The large image only shows the green and blue channel to highlight perinuclear CSPα clusters indicated by arrows. The inset is an enlarged view of the box-indicated area showing partially co-localization of CSPα-GFP with LAMP1. **b,** CSPα is partially co-localized with the LysoTracker-labeled lysosomes. *CSPα::GFP* HEK293T cells were stained with lysotracker dye (1:10,000 for 10 min at 37 °C) and imaged by live cell imaging. Nuclei were stained with Hoechst 33342. **c,** The CSPα positive perinuclear compartments are not significantly enriched with TMED10. *CSPα::GFP* U2OS cells were engineered to express TMED10 bearing an endogenously tagged mCherry at the carboxyl terminus. The cells were analyzed by live cell confocal microscopy. Nuclei were stained with Hoechst 33342. **d, e,** Co-localization of CSPα ΔNT and ΔCS with LAMP1-mCh. d, Representative confocal images of U2OS cells transfected with LAMP1-mCh together with the indicated CSPα variants. Scale bars, 5 μm. e, Quantification of the co-localization efficiency of various CSPα variants with LAMP1-mCh in individual cells in d. ****, *p* < 0.0001 by one-way ANOVA plus Dunnett’s test; n.s, not significant; *n* = 20, 16 and 25 cells, respectively.

**Figure S3, related to Figure 5.**
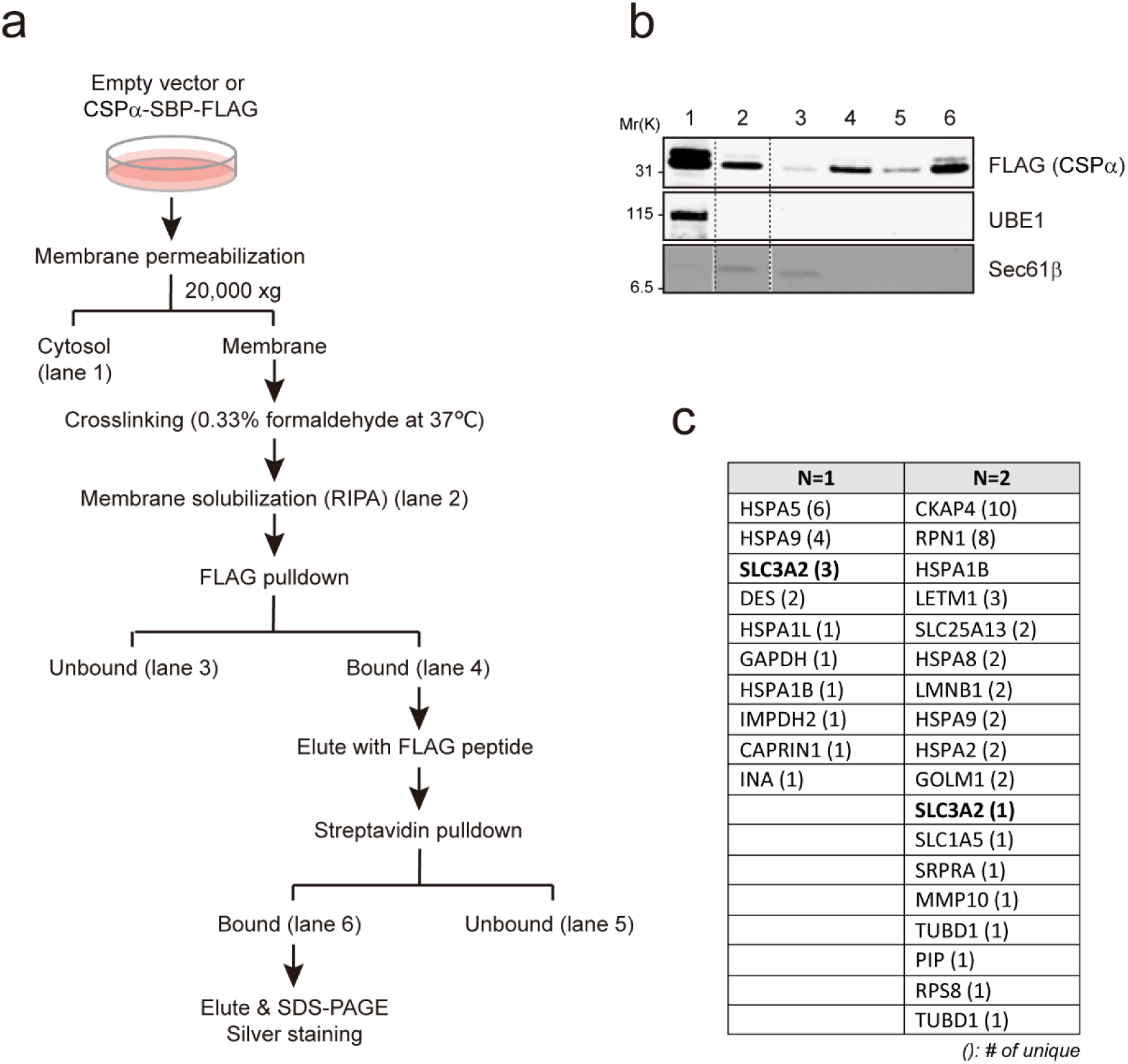
CD98hc interacts with CSPα via the LN domain and is required for the perinuclear localization of CSPα. **a,** A schematic flow chart for crosslinking and tandem affinity purification to identify CSPα ΔJ-SBP-FLAG-associated proteins. See the method section for details. **b,** Sample from the steps indicated in **a** were subjected to immunoblotting to verify the fractionation and purification procedure. **c,** Bands between 70∼100 kDa on the silver-stained gel in Fig. 5a were sectioned and analyzed by LC-MS/MS in two independent experiments. Proteins with unique peptide hits not found in the empty vector control sample are shown. The numbers in brackets indicate the number of unique peptides identified. Note that CD98hc (SLC3A2) and HSPA9 are the only hits found in both experiments.

**Figure S4, related to Figure 5.**
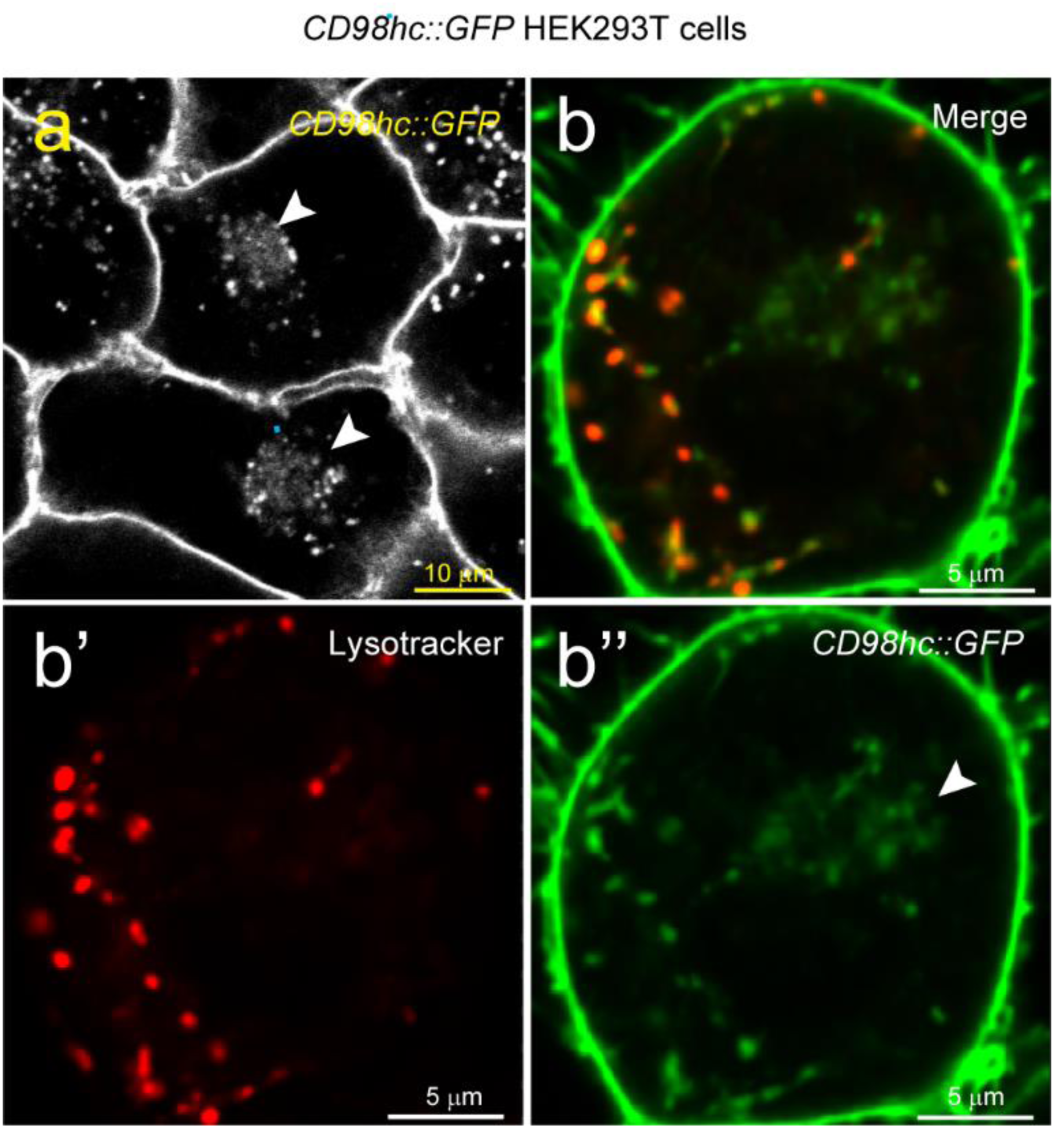
CD98hc interacts with CSPα via the LN domain and is required for the perinuclear localization of CSPα. **a,** HEK293T cells expressing CD98hc bearing an endogenously tagged GFP were imaged by live cell confocal microscopy. **b,** The cells were stained a LysoTracker dye in red for 10 min and imaged by live cell confocal microscopy. Arrowheads indicate perinuclear CD98hc clusters barely labeled by the Lysotracker dye.

**Figure S5, related to Figure 5 and 6.**
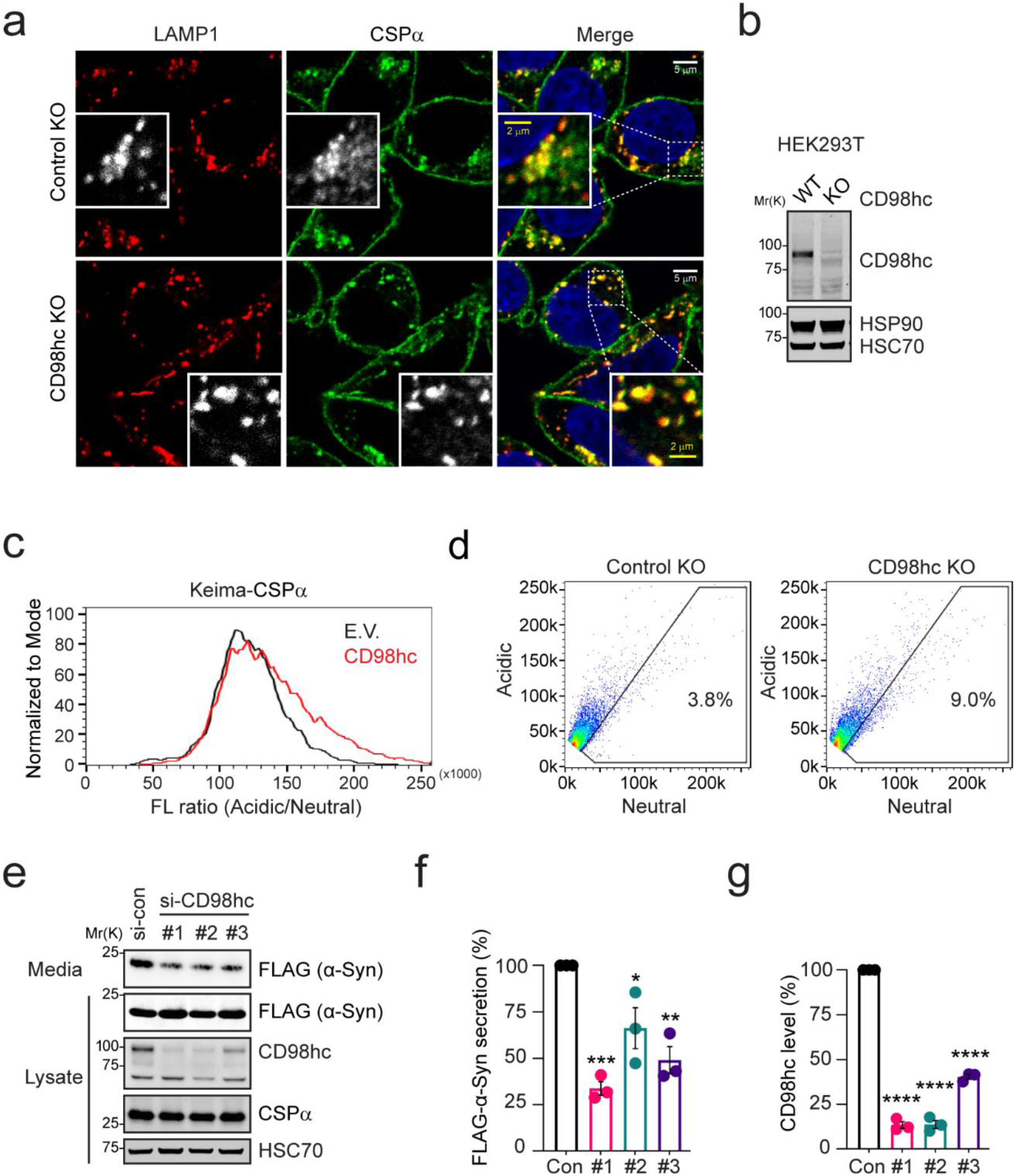
Depletion of CD98hc reduces CSPα localization to the perinuclear LAMP1 negative compartment and inhibits MAPS. **a,** CD98hc CRISPR KO or control KO HEK293T cells were double stained for endogenous LAMP1 (red) and CSPα (green) and imaged by confocal microscopy. DAPI (in blue) was used to label the nuclei. The insets show an enlarged view of the box-indicated area. **b,** The knockout of CD98hc was confirmed by immunoblotting using cell lysates prepared from a fraction of cells in a. **c,** Overexpression of CD98hc only slightly enhance the translocation of CSPα into endolysosomes. Keima-CSPα expressing HEK293T cells were transfected with either an empty vector or CD98hc-Myc-FLAG-expressing vector and analyzed by flow cytometry. The graph shows the acidic/neutral fluorescence ratio from > 10,000 cells of the Keima-positive population. **d,** Depletion of CD98hc did not significantly affect the translocation of CSPα into endolysosomes. Control KO and CD98hc KO HEK293T cells were transiently transfected with Keima-CSPα and analyzed by flow cytometry. Dot plots show a selected neutral gate, indicating that most Keima-CSPα is still translocated into endolysosomes in CD98hc KO cells. **e,** CD98hc is required for α-Syn secretion. HEK293T cells were transfected with either control siRNA or three different siRNA targeting CD98hc. 24 h post transfection, cells were re-seeded and further transfected with FLAG-human α-Synuclein. The conditioned medium (16 h) and lysates were analyzed by immunoblotting as indicated. **f,** Quantification of the α-Syn secretion from the experiments in e. error bars, mean ± s.e.m., *n* = 3 independent experiments. ***, *p* = 0.0004), \*\**p* = 0.002, \**p* = 0.0207 by one-way ANOVA plus Dunnett’s test. **g,** Quantification of the CD98hc levels from the experiments in e. error bars, mean ± s.e.m., *n* = 3 independent experiments. ****, *p* < 0.0001 by one-way ANOVA plus Dunnett’s test.

**Figure S6, related to Figure 6.**
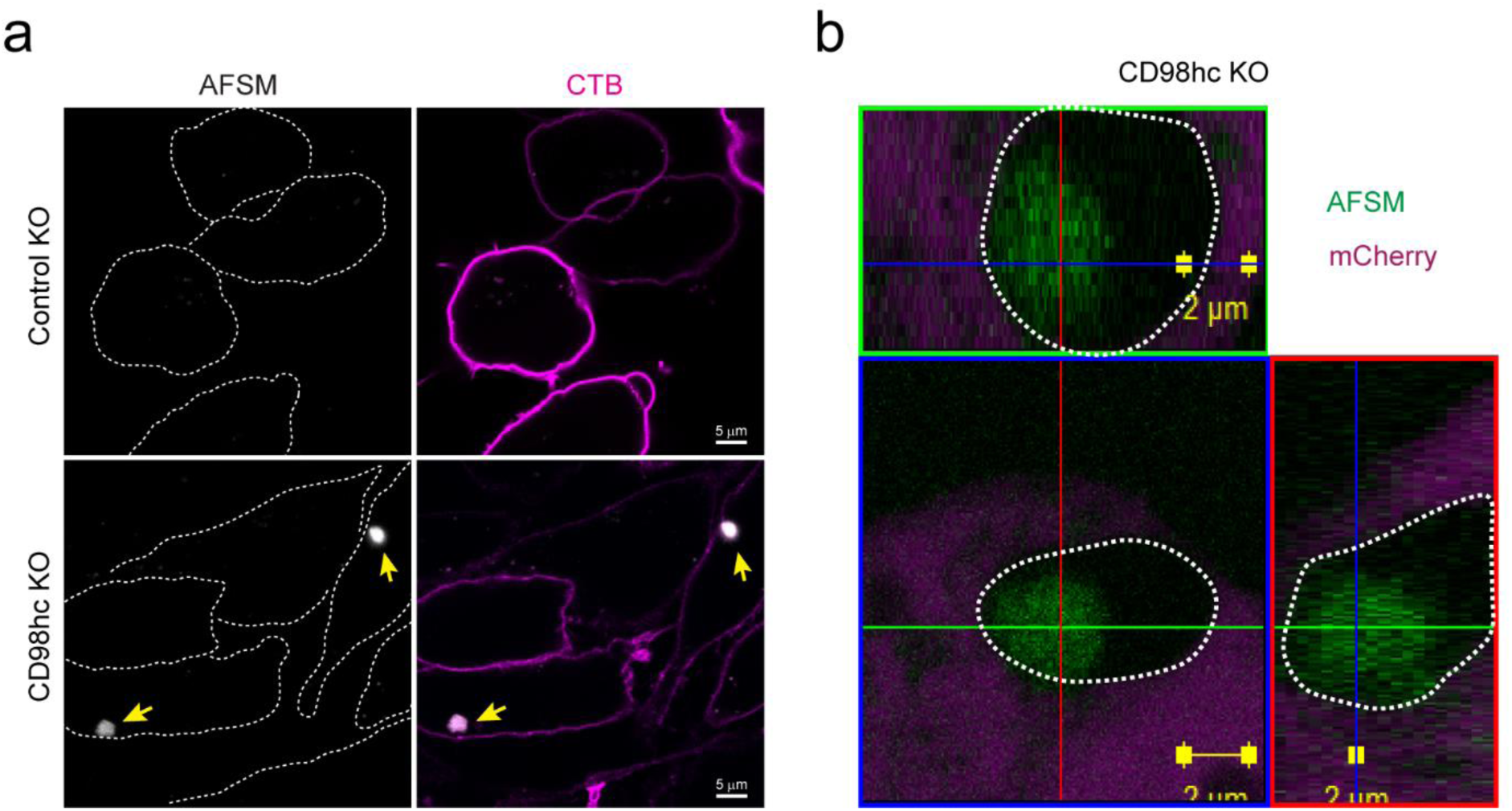
Depletion of CD98hc causes accumulation of AFSMs in cells. **a,** Control KO or CD98hc KO HEK293T cells were treated with cholera toxin subunit B (CTB) to label cell boundary. Cells were imaged by confocal microscopy. Arrows indicate AFSMs. Dashed lines show cell boundaries. **b,** Orthogonal views (x/y, x/z or y/z) of confocal z-stacks of a CD98hc KO cell transfected with mCherry (magenta). Note that the AFSM puncta (green) is localized in a cavity in the cytoplasm.

**Figure S7, related to Figure 7.**
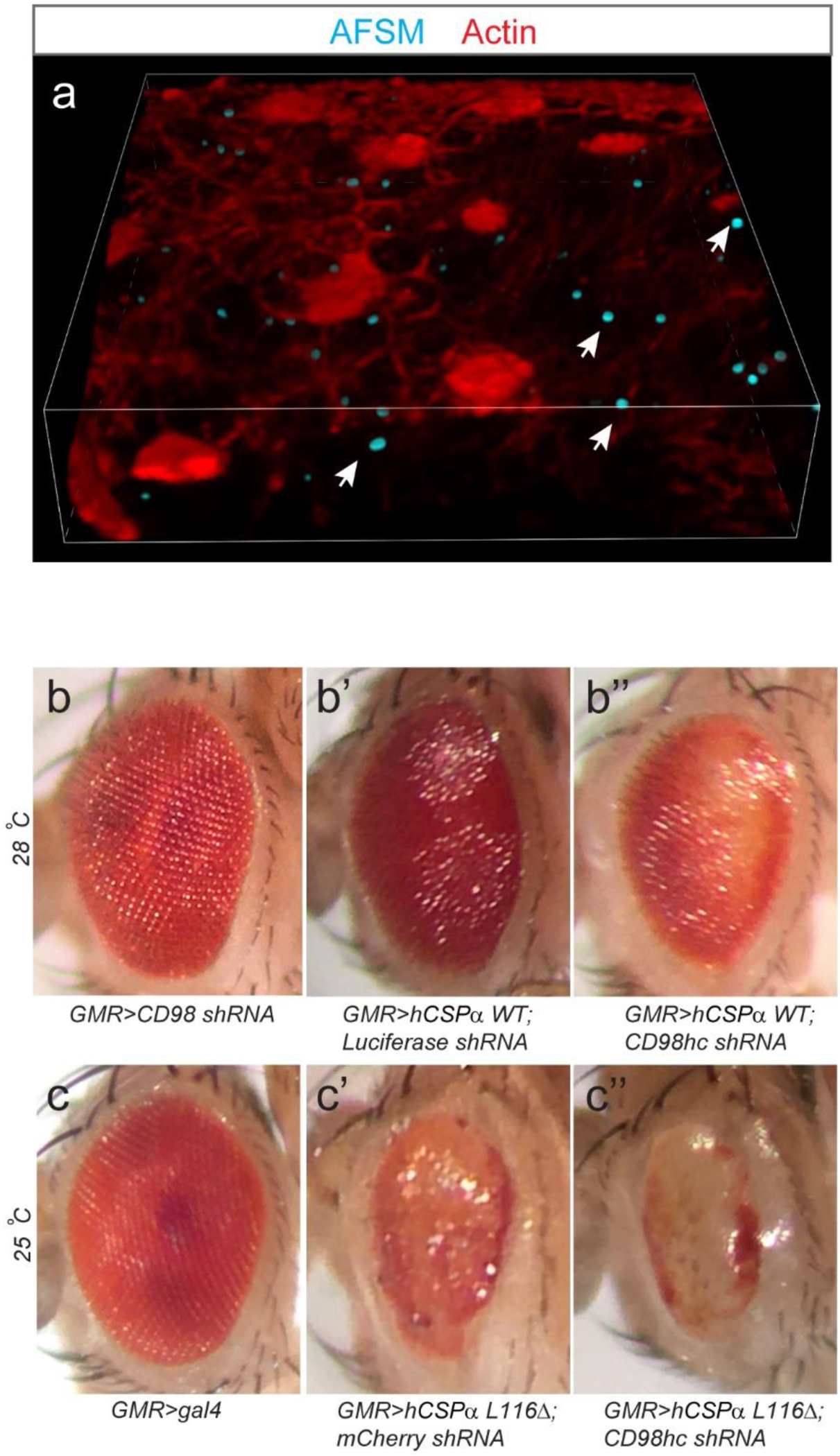
AFSM accumulation and genetic interaction between CSPα and CD98hc in a fly ANCL model. **a,** Accumulation of AFSMs in photoreceptor cells in an eye disc from a *GMR>hCSPα L116Δ* larva. Shown is a 3-D view reconstructed from confocal z-stacks of an eye disc stained by phalloidin. Arrows indicate examples of AFSMs. **b-b’’**, CD98hc knockdown enhances the rough eye phenotype associated with hCSPα WT expression at 28 °C. **c-c’’**, CD98hc knockdown enhances the rough eye phenotype associated with hCSPα L116Δ expression at 25 °C.

**Supplementary video 1**

Co-localization of endogenous CSPα and CD98hc. HEK293 cells bearing GFP and mCherry tag on CSPα and CD98hc respectively were imaged by dual-color live cell confocal microscopy. Hoechst 33342 was used for staining of nuclei.

**Supplementary video 2**

Co-localization of endogenous CSPα and lysotracker positive compartments. *CSPα::GFP* HEK293T cells were stained with a Lysotracker dye (1:10,000) and imaged in real time by a confocal microscope. Hoechst 33342 was used for staining of nuclei.

**Supplementary table 1**

Table with information on used plasmids, si-RNAs, chemicals and antibodies.

## References

1 Anderson, G. W., Goebel, H. H. & Simonati, A. Human pathology in NCL. Biochim Biophys Acta 1832, 1807–1826, doi:10.1016/j.bbadis.2012.11.014 (2013).

2 Naseri, N., Sharma, M. & Velinov, M. Autosomal dominant neuronal ceroid lipofuscinosis: Clinical features and molecular basis. Clin Genet 99, 111–118, doi:10.1111/cge.13829 (2021).

3 Haltia, M. The neuronal ceroid-lipofuscinoses: from past to present. Biochim Biophys Acta 1762, 850–856, doi:10.1016/j.bbadis.2006.06.010 (2006).

4 Cotman, S. L., Karaa, A., Staropoli, J. F. & Sims, K. B. Neuronal ceroid lipofuscinosis: impact of recent genetic advances and expansion of the clinicopathologic spectrum. Curr Neurol Neurosci Rep 13, 366, doi:10.1007/s11910-013-0366-z (2013).

5 Specchio, N. et al. Neuronal Ceroid Lipofuscinosis: Potential for Targeted Therapy. Drugs, doi:10.1007/s40265-020-01440-7 (2020).

6 Mole, S. E. & Cotman, S. L. Genetics of the neuronal ceroid lipofuscinoses (Batten disease). Biochim Biophys Acta 1852, 2237–2241, doi:10.1016/j.bbadis.2015.05.011 (2015).

7 Bajaj, L. et al. A CLN6-CLN8 complex recruits lysosomal enzymes at the ER for Golgi transfer. J Clin Invest 130, 4118–4132, doi:10.1172/JCI130955 (2020).

8 di Ronza, A. et al. CLN8 is an endoplasmic reticulum cargo receptor that regulates lysosome biogenesis. Nat Cell Biol 20, 1370–1377, doi:10.1038/s41556-018-0228-7 (2018).

9 Chamberlain, L. H. & Burgoyne, R. D. Cysteine-string protein: the chaperone at the synapse. J Neurochem 74, 1781–1789, doi:10.1046/j.1471-4159.2000.0741781.x (2000).

10 Greaves, J., Salaun, C., Fukata, Y., Fukata, M. & Chamberlain, L. H. Palmitoylation and membrane interactions of the neuroprotective chaperone cysteine-string protein. J Biol Chem 283, 25014–25026, doi:10.1074/jbc.M802140200 (2008).

11 Greaves, J. & Chamberlain, L. H. Dual role of the cysteine-string domain in membrane binding and palmitoylation-dependent sorting of the molecular chaperone cysteine-string protein. Mol Biol Cell 17, 4748–4759, doi:10.1091/mbc.e06-03-0183 (2006).

12 Xu, Y. et al. DNAJC5 facilitates USP19-dependent unconventional secretion of misfolded cytosolic proteins. Cell Discov 4, 11, doi:10.1038/s41421-018-0012-7 (2018).

13 Benitez, B. A. & Sands, M. S. Primary fibroblasts from CSPalpha mutation carriers recapitulate hallmarks of the adult onset neuronal ceroid lipofuscinosis. Sci Rep 7, 6332, doi:10.1038/s41598-017-06710-1 (2017).

14 Zinsmaier, K. E. et al. A cysteine-string protein is expressed in retina and brain of Drosophila. J Neurogenet 7, 15–29, doi:10.3109/01677069009084150 (1990).

15 Ohyama, T. et al. Huntingtin-interacting protein 14, a palmitoyl transferase required for exocytosis and targeting of CSP to synaptic vesicles. J Cell Biol 179, 1481–1496, doi:10.1083/jcb.200710061 (2007).

16 Tobaben, S. et al. A trimeric protein complex functions as a synaptic chaperone machine. Neuron 31, 987–999, doi:10.1016/s0896-6273(01)00427-5 (2001).

17 Braun, J. E., Wilbanks, S. M. & Scheller, R. H. The cysteine string secretory vesicle protein activates Hsc70 ATPase. J Biol Chem 271, 25989–25993, doi:10.1074/jbc.271.42.25989 (1996).

18 Chamberlain, L. H. & Burgoyne, R. D. Activation of the ATPase activity of heat-shock proteins Hsc70/Hsp70 by cysteine-string protein. Biochem J 322 **(Pt** **3**), 853–858, doi:10.1042/bj3220853 (1997).

19 Sharma, M. et al. CSPalpha knockout causes neurodegeneration by impairing SNAP-25 function. EMBO J 31, 829–841, doi:10.1038/emboj.2011.467 (2012).

20 Sharma, M., Burre, J. & Sudhof, T. C. CSPalpha promotes SNARE-complex assembly by chaperoning SNAP-25 during synaptic activity. Nat Cell Biol 13, 30–39, doi:10.1038/ncb2131 (2011).

21 Ranjan, R., Bronk, P. & Zinsmaier, K. E. Cysteine string protein is required for calcium secretion coupling of evoked neurotransmission in drosophila but not for vesicle recycling. J Neurosci 18, 956–964 (1998).

22 Miller, L. C. et al. Cysteine string protein (CSP) inhibition of N-type calcium channels is blocked by mutant huntingtin. J Biol Chem 278, 53072–53081, doi:10.1074/jbc.M306230200 (2003).

23 Weng, N. et al. Functional role of J domain of cysteine string protein in Ca2+-dependent secretion from acinar cells. Am J Physiol Gastrointest Liver Physiol 296, G1030–1039, doi:10.1152/ajpgi.90592.2008 (2009).

24 Bronk, P. et al. The multiple functions of cysteine-string protein analyzed at Drosophila nerve terminals. J Neurosci 25, 2204–2214, doi:10.1523/JNEUROSCI.3610-04.2005 (2005).

25 Bronk, P. et al. Drosophila Hsc70-4 is critical for neurotransmitter exocytosis in vivo. Neuron 30, 475–488, doi:10.1016/s0896-6273(01)00292-6 (2001).

26 Nie, Z. et al. Overexpression of cysteine-string proteins in Drosophila reveals interactions with syntaxin. J Neurosci 19, 10270–10279 (1999).

27 Evans, G. J. & Morgan, A. Phosphorylation-dependent interaction of the synaptic vesicle proteins cysteine string protein and synaptotagmin I. Biochem J 364, 343–347, doi:10.1042/BJ20020123 (2002).

28 Wu, M. N. et al. Syntaxin 1A interacts with multiple exocytic proteins to regulate neurotransmitter release in vivo. Neuron 23, 593–605, doi:10.1016/s0896-6273(00)80811-9 (1999).

29 Fernandez-Chacon, R. et al. The synaptic vesicle protein CSP alpha prevents presynaptic degeneration. Neuron 42, 237–251, doi:10.1016/s0896-6273(04)00190-4 (2004).

30 Gorenberg, E. L. & Chandra, S. S. The Role of Co-chaperones in Synaptic Proteostasis and Neurodegenerative Disease. Front Neurosci 11, 248, doi:10.3389/fnins.2017.00248 (2017).

31 Donnelier, J. & Braun, J. E. CSPalpha-chaperoning presynaptic proteins. Front Cell Neurosci 8, 116, doi:10.3389/fncel.2014.00116 (2014).

32 Lee, J. G., Takahama, S., Zhang, G., Tomarev, S. I. & Ye, Y. Unconventional secretion of misfolded proteins promotes adaptation to proteasome dysfunction in mammalian cells. Nat Cell Biol 18, 765–776, doi:10.1038/ncb3372 (2016).

33 Fontaine, S. N. et al. DnaJ/Hsc70 chaperone complexes control the extracellular release of neurodegenerative-associated proteins. EMBO J 35, 1537–1549, doi:10.15252/embj.201593489 (2016).

34 Noskova, L. et al. Mutations in DNAJC5, encoding cysteine-string protein alpha, cause autosomal-dominant adult-onset neuronal ceroid lipofuscinosis. Am J Hum Genet 89, 241–252, doi:10.1016/j.ajhg.2011.07.003 (2011).

35 Benitez, B. A. et al. Exome-sequencing confirms DNAJC5 mutations as cause of adult neuronal ceroid-lipofuscinosis. PLoS One 6, e26741, doi:10.1371/journal.pone.0026741 (2011).

36 Cadieux-Dion, M. et al. Recurrent mutations in DNAJC5 cause autosomal dominant Kufs disease. Clin Genet 83, 571–575, doi:10.1111/cge.12020 (2013).

37 Diez-Ardanuy, C., Greaves, J., Munro, K. R., Tomkinson, N. C. & Chamberlain, L. H. A cluster of palmitoylated cysteines are essential for aggregation of cysteine-string protein mutants that cause neuronal ceroid lipofuscinosis. Sci Rep 7, 10, doi:10.1038/s41598-017-00036-8 (2017).

38 Imler, E. et al. A Drosophila model of neuronal ceroid lipofuscinosis CLN4 reveals a hypermorphic gain of function mechanism. Elife 8, doi:10.7554/eLife.46607 (2019).

39 Naseri, N. N. et al. Aggregation of mutant cysteine string protein-alpha via Fe-S cluster binding is mitigated by iron chelators. Nat Struct Mol Biol 27, 192–201, doi:10.1038/s41594-020-0375-y (2020).

40 Violot, S., Carpentier, P., Blanchoin, L. & Bourgeois, D. Reverse pH-dependence of chromophore protonation explains the large Stokes shift of the red fluorescent protein mKeima. J Am Chem Soc 131, 10356–10357, doi:10.1021/ja903695n (2009).

41 Katayama, H., Kogure, T., Mizushima, N., Yoshimori, T. & Miyawaki, A. A sensitive and quantitative technique for detecting autophagic events based on lysosomal delivery. Chem Biol 18, 1042–1052, doi:10.1016/j.chembiol.2011.05.013 (2011).

42 Polymeropoulos, M. H. et al. Mutation in the alpha-synuclein gene identified in families with Parkinson’s disease. Science 276, 2045–2047, doi:10.1126/science.276.5321.2045 (1997).

43 Sardana, R. & Emr, S. D. Membrane Protein Quality Control Mechanisms in the Endo-Lysosome System. Trends Cell Biol 31, 269–283, doi:10.1016/j.tcb.2020.11.011 (2021).

44 Sahu, R. et al. Microautophagy of cytosolic proteins by late endosomes. Dev Cell 20, 131–139, doi:10.1016/j.devcel.2010.12.003 (2011).

45 Feng, Y., He, D., Yao, Z. & Klionsky, D. J. The machinery of macroautophagy. Cell Res 24, 24–41, doi:10.1038/cr.2013.168 (2014).

46 Lee, C., Lamech, L., Johns, E. & Overholtzer, M. Selective Lysosome Membrane Turnover Is Induced by Nutrient Starvation. Dev Cell 55, 289–297 e284, doi:10.1016/j.devcel.2020.08.008 (2020).

47 Mesquita, A., Glenn, J. & Jenny, A. Differential activation of eMI by distinct forms of cellular stress. Autophagy, 1–13, doi:10.1080/15548627.2020.1783833 (2020).

48 Rosell, A. et al. Structural bases for the interaction and stabilization of the human amino acid transporter LAT2 with its ancillary protein 4F2hc. Proc Natl Acad Sci U S A 111, 2966–2971, doi:10.1073/pnas.1323779111 (2014).

49 Zhang, M. et al. A Translocation Pathway for Vesicle-Mediated Unconventional Protein Secretion. Cell 181, 637–652 e615, doi:10.1016/j.cell.2020.03.031 (2020).

50 Tyynela, J., Palmer, D. N., Baumann, M. & Haltia, M. Storage of saposins A and D in infantile neuronal ceroid-lipofuscinosis. FEBS Lett 330, 8–12, doi:10.1016/0014-5793(93)80908-d (1993).

51 Lojewski, X. et al. Human iPSC models of neuronal ceroid lipofuscinosis capture distinct effects of TPP1 and CLN3 mutations on the endocytic pathway. Hum Mol Genet 23, 2005–2022, doi:10.1093/hmg/ddt596 (2014).

52 Greaves, J. et al. Palmitoylation-induced aggregation of cysteine-string protein mutants that cause neuronal ceroid lipofuscinosis. J Biol Chem 287, 37330–37339, doi:10.1074/jbc.M112.389098 (2012).

53 Milkereit, R. et al. LAPTM4b recruits the LAT1-4F2hc Leu transporter to lysosomes and promotes mTORC1 activation. Nat Commun 6, 7250, doi:10.1038/ncomms8250 (2015).

54 Bruns, C., McCaffery, J. M., Curwin, A. J., Duran, J. M. & Malhotra, V. Biogenesis of a novel compartment for autophagosome-mediated unconventional protein secretion. J Cell Biol 195, 979–992, doi:10.1083/jcb.201106098 (2011).

55 Malhotra, V. Unconventional protein secretion: an evolving mechanism. EMBO J 32, 1660–1664, doi:10.1038/emboj.2013.104 (2013).

56 Patel, P., Prescott, G. R., Burgoyne, R. D., Lian, L. Y. & Morgan, A. Phosphorylation of Cysteine String Protein Triggers a Major Conformational Switch. Structure 24, 1380–1386, doi:10.1016/j.str.2016.06.009 (2016).

57 Henderson, M. X. et al. Neuronal ceroid lipofuscinosis with DNAJC5/CSPalpha mutation has PPT1 pathology and exhibit aberrant protein palmitoylation. Acta Neuropathol 131, 621–637, doi:10.1007/s00401-015-1512-2 (2016).

58 Ran, F. A. et al. Genome engineering using the CRISPR-Cas9 system. Nat Protoc 8, 2281–2308, doi:10.1038/nprot.2013.143 (2013).

59 Shalem, O. et al. Genome-scale CRISPR-Cas9 knockout screening in human cells. Science 343, 84–87, doi:10.1126/science.1247005 (2014).

60 Fueller, J. et al. CRISPR-Cas12a-assisted PCR tagging of mammalian genes. J Cell Biol 219, doi:10.1083/jcb.201910210 (2020).

